# Activity-dependent synthesis of Emerin gates neuronal plasticity by regulating proteostasis

**DOI:** 10.1101/2024.06.30.600712

**Authors:** Yi Xie, Ruoxi Wang, Daniel B. McClatchy, Yuanhui Ma, Jolene Diedrich, Manuel Sanchez-Alavez, Michael Petrascheck, John R. Yates, Hollis T. Cline

## Abstract

Neurons dynamically regulate their proteome in response to sensory input, a key process underlying experience-dependent plasticity. We characterized the visual experience-dependent nascent proteome within a brief, defined time window after stimulation using an optimized metabolic labeling approach. Visual experience induced cell type-specific and age-dependent alterations in the nascent proteome, including proteostasis-related processes. We identified Emerin as the top activity-induced candidate plasticity protein and demonstrated that its rapid activity-induced synthesis is transcription-independent. In contrast to its nuclear localization and function in myocytes, activity-induced neuronal Emerin is abundant in the endoplasmic reticulum and broadly inhibits protein synthesis, including translation regulators and synaptic proteins. Downregulating Emerin shifted the dendritic spine population from predominantly mushroom morphology to filopodia and decreased network connectivity. In mice, decreased Emerin reduced visual response magnitude and impaired visual information processing. Our findings support an experience-dependent feed-forward role for Emerin in temporally gating neuronal plasticity by negatively regulating translation.

## Introduction

Sensory experience exquisitely regulates animals’ sensory function and behavior. Visual experience guides the structural and functional organization of the visual system during development [1–3], and continues to instruct circuit rewiring in adults to fine-tune neuronal activity in response to task-relevant environmental cues [4–6]. These activity-dependent processes allow continuous integration of past experiences into neural networks by modifying the physiological properties [7, 8] and synaptic connections [9–11] of distinct neuronal cell types [12–14]. Experience-dependent neuronal plasticity is not only crucial for survival behaviors such as prey capture [15, 16] and depth perception [17], but is also an underlying mechanism for cognitive functions such as learning and memory [18, 19]. A comprehensive understanding of the molecular programs orchestrating experience-dependent plasticity is key to deciphering its mechanisms and to reversing pathology in disease states.

Proteins are crucial substrates underlying neuronal function and plasticity. Sensory input greatly influences the cellular proteomic composition, which then mediates structural and functional alterations [20, 21]. Changes in the proteome can be driven by gene transcription [22–24], as well as a myriad of transcription-independent processes, giving rise to multiple layers of spatial and temporal complexity of the proteome [25]. These processes, falling under the umbrella of proteostasis, include translational control by translational factors [26], local protein synthesis from readily available mRNAs [27], RNA-binding proteins buffering the translation of target mRNAs [28, 29], protein trafficking [30, 31] and proteosome-dependent degradation [32, 33]. Proteostasis is a crucial component for neuronal function and plasticity in both health and disease [34–36], however, it is not well understood how proteostasis regulation is recruited by sensory input and how different aspects of proteostasis are temporally coordinated in response to neuronal activity.

To explore these questions, we evaluated the rapid dynamics of the visual experience-dependent nascent proteome in excitatory and inhibitory neurons in young mice during the visual critical period and in adult mice. We used metabolic labeling to identify the newly-synthesized proteome [37] with improved temporal resolution and incorporated_direct MS/MS <u>d</u>etection of <u>bi</u>otin-tag (DiDBiT) on labeled peptides [38] into the tandem mass spectrometry (MS/MS) pipeline, increasing the detection efficiency and accuracy. We found an activity-dependent enrichment in proteins related to proteostasis across neuronal cell types and ages, suggesting its broad participation in experience-dependent plasticity. The top experience-dependent candidate plasticity protein (CPP), Emerin, is rapidly synthesized independent of transcription. In contrast to its nuclear envelope localization and functions in myocytes, activity-induced nascent Emerin in neurons is abundant in the ER and broadly reduces protein synthesis, suggesting Emerin’s multifunctionality arises from its activity-dependent regulation and tissue-specific localization. Downregulating Emerin shifts the population of dendritic spines from mature mushroom morphology to immature filopodial morphology and reduces synaptic connectivity. In mice, decreasing Emerin reduced the magnitude of visually evoked responses and impaired visual information processing. Our data suggest activity-induced Emerin functions in a feed-forward role regulating neuronal proteostasis and activity-dependent plasticity.

## Results

### An optimized proteomic pipeline targeting acute cell type-specific nascent proteome

Analyzing rapid experience-dependent changes in the nascent proteome with cell-type resolution could offer mechanistic insights into neuronal plasticity. <u>B</u>io-<u>o</u>rthogonal <u>n</u>on<u>c</u>anonical <u>a</u>mino <u>a</u>cid tagging (BONCAT) provides a cell type-specific approach to metabolically label the nascent proteome using the noncanonical amino acid azidonorleucine (ANL) and a mouse line expressing <u>m</u>utant <u>met</u>hionyl-t<u>R</u>NA <u>s</u>ynthetase (mMetRS^KI/KI^) crossed to cre drivers with cell-type specificity (Figure S1A). However, shortening the labeling period to improve temporal sensitivity poses a great challenge for proteomics analysis due to the small amounts of labeled material. To enable a temporally-sensitive, cell type-specific nascent proteome screen, we optimized the proteomics pipeline by maximizing ANL labeling efficiency and MS/MS detection sensitivity. We previously showed that the concentration of free ANL in the brain reached a plateau between 1-4h after a single intraperitoneal (i.p.) injection and was undetectable after 8h [39]. To identify the in vivo kinetics of ANL protein labeling in cortex, we collected cortical tissue at 2, 4, 8, 16 and 24h after i.p. ANL injections, biotinylated ANL-labeled proteins in protein lysates with biotin-alkyne click chemistry and quantified biotin-labeled protein in western blots. Biotinylated protein levels peaked at 8h after ANL i.p. injection (Figure S1B), indicating an optimal timepoint to measure activity-induced nascent proteins. Moreover, we showed that homozygous mMetRS^KI/KI^ improved labeling efficiency ∼2-fold compared to their heterozygous littermates in both EMX^cre^+ (Figure S1C) and vGat^cre^+ (Figure S1D) neurons. These data allowed us to shorten the protein labeling window to 8-10h after ANL delivery, without compromising the subsequent proteome coverage.

Biotinylation of ANL-labeled proteins allows the enrichment of the labeled proteome with neutravidin beads. However, this process is often complicated by non-specific binding to beads, especially in the case of cell type-specific analysis of rapid proteome dynamics where the unlabeled protein pool is significantly larger than the targeted protein pool. To avoid false-positive hits and increase MS/MS detection sensitivity, we adopted the BONCAT-DiDBiT method which identifies the biotin tag on methionine with MS/MS and distinguishes true hits from contaminants [39]. Experimental and control samples were duplexed using stable heavy/light-isotope labeled biotin-alkyne, which avoids introducing sample variability during processing and thus increases sensitivity of the assay (Figure S1E). Together, this BONCAT-DiDBiT-MS/MS pipeline allowed us to target rapid experience-dependent, cell type-specific changes in the nascent proteome.

### Visual experience-induced nascent proteomes reveal a common theme of proteostasis regulation across cell types and ages

To identify visual experience-dependent nascent proteome dynamics in excitatory and inhibitory visual cortical neurons, we dark-housed male and female critical period (P28) and adult (P56) mice for 3 days and delivered ANL by i.p. injection immediately before either 4h of visual experience (VE) or 4h in total darkness (DK). We collected the visual cortex 6h later and prepared samples for BONCAT and DiDBiT (Figure 1A). VE and DK samples were labeled with heavy-or light-biotin-alkyne, respectively, by click chemistry and duplexed during sample processing and MS/MS detection. We tested the efficacy of our DK-VE treatment on visual cortex activity by comparing cFos labeling in the visual cortex of DK and VE mice. Consistent with previous studies showing that similar DK/VE paradigm potently induces activity-dependent genes [12, 40], our VE treatment induced robust cFos expression compared to the DK group (Figure 1A), which validates the use of this VE paradigm to assess the activity-dependent nascent proteome. Then, we evaluated both excitatory and inhibitory VE-dependent nascent proteomes using the EMX^cre^-mMetRS^KI/KI^ and vGat^cre^-mMetRS^KI/KI^ mice (Figure 1A). We detected 2489, 1995, 1944 and 1418 biotinylated nascent proteins in P28 excitatory neurons (EX28), P28 inhibitory neurons (IN28), P56 excitatory neurons (EX56) and P56 inhibitory neurons (IN56), respectively, of which 278 (EX28), 231 (IN28), 364 (EX56) and 253 (IN56) proteins were significantly regulated by VE (Figure 1B, Table S1). These VE-induced proteins respond to sensory input at the proteome level and may play a role in regulating various aspects of activity-dependent plasticity. Therefore, we named them candidate plasticity proteins (CPPs). The number of CPPs constitutes >10% of the total biotinylated proteins detected, demonstrating the capability of our pipeline to detect the rapid dynamics of the nascent proteome. Furthermore, VE resulted in significant increases and decreases CPPs. Specifically, during the critical period, VE significantly increased 214 CPPs and significantly decreased 64 CPPS in EX28 neurons, and significantly increased 140 CPPs and significantly decreased 91 CPPS in IN28 neurons. In mature animals, VE significantly increased 228 CPPs and significantly decreased 136 CPPS in EX56 neurons, and significantly increased 200 CPPs and significantly decreased 53 CPPS in IN56 neurons.

**Figure 1.**
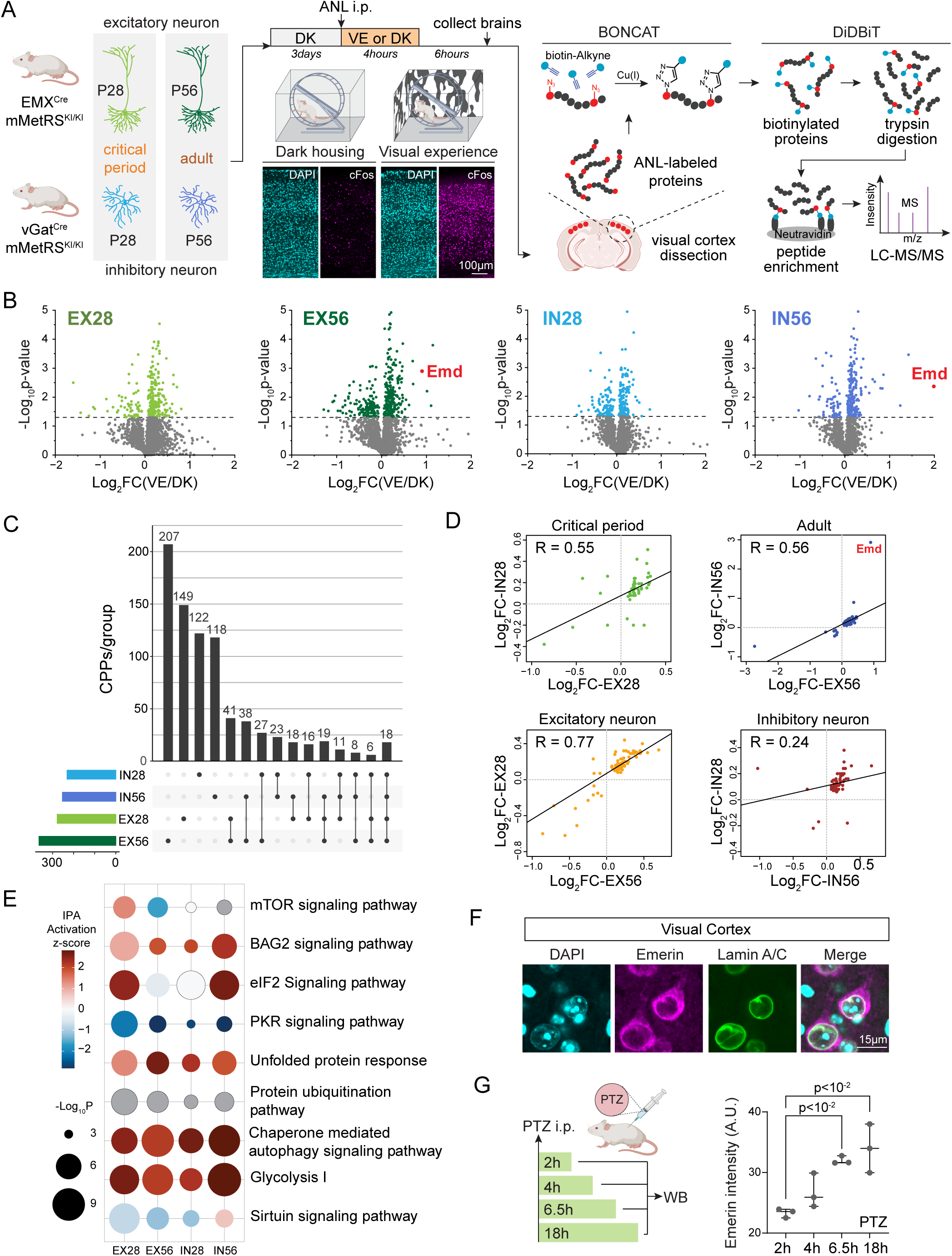
Cell type-specific proteomics characterize visual experience-dependent nascent protein dynamics. (A) Proteomics workflow. EMX^cre^-mMetRS^KI/KI^ and vGat^cre^-mMetRS^KI/KI^ mice at P28 and P56 were dark-housed, injected with ANL and then either exposed to visual experience (VE) or dark (DK). Visual cortices were dissected, processed and analyzed following BONCAT-DiDBiT pipeline. (B-D) Proteomics datasets. (B) Volcano plots visualize all proteins identified in each condition. n=6 biological replicates for each dataset. Colored dots, candidate plasticity proteins (CPPs, p<0.05). (C) Upset plot showing intersection sizes of CPPs under different experimental conditions. (D) Correlation of CPPs shared between two conditions with the same age or cell type. R, Pearson correlation coefficient. (E) Significantly regulated pathways shared by all conditions. IPA activation z-score indicates IPA-predicted pathway activation (red) or inhibition (blue). Grey, prediction not available. (F) Emerin immunostaining in mouse primary visual cortex. DAPI, nuclear marker. Lamin A/C in the inner nuclear membrane. (G) PTZ-induced Emerin expression in visual cortex. Left, experimental workflow. Right, western blot quantification of Emerin signal at 2h, 4h, 6.5h and 18h after PTZ i.p. injection, normalized to ponceau. n=3, one-way ANOVA with Sidak multiple comparison (all timepoints compared to 2h). Lines indicate median and 95% CI.

The majority of CPPs (596/821) are unique to each condition, suggesting considerable diversity of experience-dependent regulatory mechanisms across cell types and ages (Figure 1C). Correlation analysis of the shared CPPs by the same cell type or age showed a mild positive correlation in each comparison (R ∈ [0.24-0.77]) (Figure 1D), suggesting both similarities and differences in CPPs across conditions. Then, we asked whether VE-dependent regulatory elements were shared by all cell types and ages. Using Ingenuity Pathway Analysis (IPA), we extracted significant VE-regulated pathways in all tested conditions (p < 0.001). Interestingly, apart from metabolic pathways, the majority of pathways in common across conditions concern various aspects of proteostasis regulation, including signal transduction pathways upstream of translation (mTOR signaling pathway, BAG2-MAPK signaling pathway), protein translational control (eIF2 signaling pathway, PKR signaling pathway), ER processing of protein (unfolded protein response) and protein degradation pathways (protein ubiquitination pathway and chaperone-mediated autophagy signaling pathway) (Figure 1E, Table S2). Pathway analysis using Metascape also revealed a broad activity-induced regulation of proteostasis, which includes shared pathway clusters related to the terms eukaryotic translation initiation (Figure S2A), vesicle-mediated protein transportation (Figure S2B), protein processing in the ER (Figure S2C) and protein localization regulation (Figure S2D, Table S3). Overall, these analyses identified diverse activity-dependent protein regulatory programs as well as a common theme of proteostasis regulation.

### Emerin is an activity-regulated protein

Emerin emerged at the top of the CPP list with the highest VE-induced expression fold change. Emerin is expressed in every layer of visual cortex (Figure S3A) and is upregulated by VE in both excitatory and inhibitory neurons in adult visual cortex (Figure S3B), however, its function in neurons is unclear. Our bioinformatic analysis suggests that Emerin might physically interact with other CPPs in a network regulating protein localization (GO:0008104, Figure S2D). As a genetic risk factor for Emery-Dreifuss muscular dystrophy, studies on Emerin have largely focused on skeletal and cardiac myocytes where it is largely located in the inner nuclear membrane through high-affinity binding with Lamin A [41–43] and modulates chromatin structure and transcription factor signaling by interacting with other nuclear proteins [44, 45]. Emerin also affects nuclear structure by increasing actin polymerization in the nuclear cortical network [46], a function that Emerin may also play in regulating cytosolic actin [41]. In neurons and other cells lacking Lamin A, Emerin is located in the nuclear envelope (NE) and the plasma membrane, and is enriched in the ER [41, 42, 47, 48], the major cellular compartment regulating protein synthesis and post-translational processes. Emerin has distinct functional domains and multiple interacting partners, suggesting that it may have diverse functions depending on its subcellular location [49, 50]. Emerin immunolabeling in mouse visual cortical sections (Figure 1F) and primary neuronal cultures (Figure 2B) indicates a predominant extranuclear cytoplasmic localization, consistent with distribution in the ER (Figure S3D). Emerin’s distinct neuronal localization pattern and activity-dependent expression prompted us to explore its function in activity-dependent neuronal plasticity.

**Figure 2.**
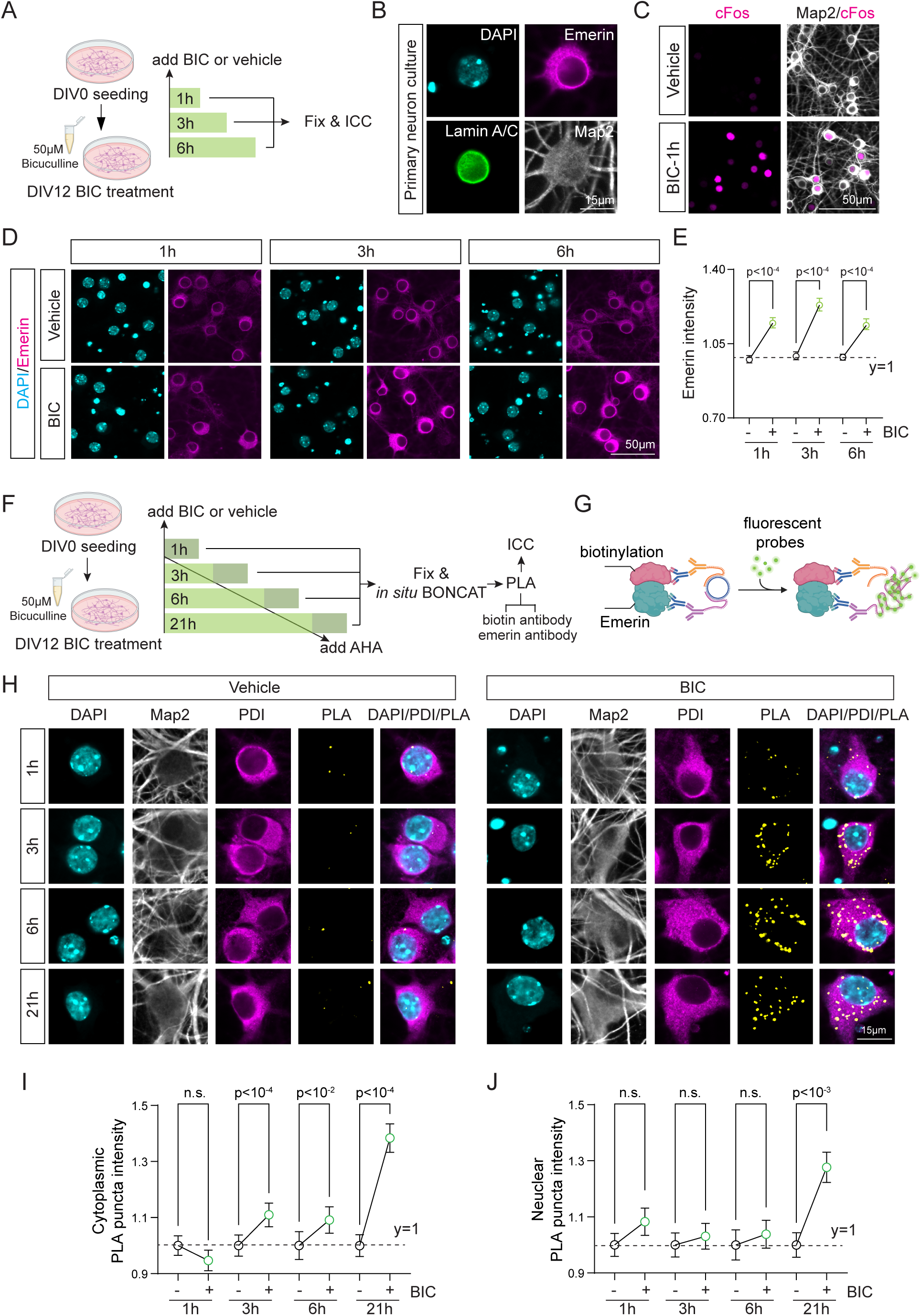
Emerin is an activity-regulated protein. (A-E) Timecourse of activity-induced Emerin expression. (A) Experimental workflow. DIV12 primary cortical neuronal cultures were treated with bicuculline (BIC) for 1h, 3h and 6h, then fixed and immunolabeled. (B) Emerin, Lamin A/C and Map2 immunolabeling. (C) cFos immunolabeling 1h after BIC or vehicle. (D) Representative images of Emerin immunolabeling 1h, 3h and 6h after BIC or vehicle treatment. (E) Quantification of Emerin labeling. BIC groups are normalized to the corresponding vehicle controls. n=1000-1500 individual neurons from 4 wells of 2 independent culture preparations. Kruskal-Wallis test with Dunn’s multiple comparison (compare BIC to the corresponding vehicle). Dots and lines indicate median and 95% CI. (F-J) Timecourse of activity-induced Emerin synthesis using in situ BONCAT-PLA. (F) Experimental workflow. DIV12 neuronal cultures were treated with BIC for 1h, 3h, 6h and 21h. AHA was added during the last hour of BIC treatment. Samples were processed with *in situ* BONCAT-PLA. (G) Schematic of *in situ* BONCAT-PLA. Antibodies recognizing biotin and Emerin detect newly-synthesized Emerin. (H) Representative images showing DAPI, Map2, PDI, an ER marker, and BONCAT-Emerin-PLA puncta following BIC/vehicle treatment at each timepoint. (I) Quantification of cytoplasmic PLA intensity. (J) Quantification of nuclear PLA intensity. Nucleus was defined by DAPI outline. BIC groups are normalized to the corresponding vehicle controls. n=1500-2500 individual neurons from 6 wells of 2 independent culture preparations. Kruskal-Wallis test with Dunn’s multiple comparison (compare BIC to the corresponding vehicle). Dots and lines indicate median and 95% CI. n.s., not significant.

We first examined how activity regulates Emerin expression. Western blot analysis independently validated the MS/MS proteomics finding that VE increased Emerin expression in mouse visual cortex (Figure S3C). Then, we asked if this VE-dependent increase in expression generalizes to a neuronal activity-dependent regulation, using i.p. injection of the GABA_A_ receptor antagonist, pentylenetetrazol (PTZ), to increase brain activity. We collected visual cortex tissue at 2, 4, 6.5 and 18h after PTZ treatment and quantified Emerin expression by western blot. PTZ induced a sustained increase in Emerin expression in mice (Figure 1G), indicating that disinhibition-induced global neuronal activation upregulated Emerin expression as well. Similarly, exposing mouse primary neuronal cultures to bicuculline, another GABA_A_ receptor antagonist, increased Emerin protein expression as quickly as 1h after treatment, similar to cFos (Figure 2A, 2C-E). Following the acute induction with PTZ or bicuculline, Emerin level remained elevated for hours after stimulation (Figure 1G, right, Figure 2D,E), suggesting that induction of Emerin might have a prolonged effect on regulating activity-dependent plasticity.

To examine the subcellular localization of activity-induced Emerin, we labeled newly-synthesized proteins by exposing cultures to AHA during the last hour of bicuculline treatments and detected the newly-synthesized Emerin using *in situ* BONCAT and proximity ligation assay (*in situ* BONCAT-PLA) (Figure 2F-G). We measured newly-synthesized Emerin in the nucleus and the extranuclear cytoplasmic compartments and found a significant accumulation of newly-synthesized Emerin puncta in the cytoplasmic compartment 3h, 6h and 21h after bicuculline stimulation (Figure 2H-I). The colocalization of these puncta with the ER marker, protein disulfide isomerase (PDI), suggests that activity-induced nascent Emerin is localized to the ER, consistent with its post-translational insertion into the ER membrane [41, 42]. By contrast, we did not detect significant changes in Emerin puncta in the nucleus until 21 hours after stimulation (Figure 2J). Together, these results characterize the activity-dependence of Emerin expression and suggest activity-induced Emerin is located at or near the ER.

### Acute activity-induced Emerin expression is driven directly by translation

To further probe the mechanism underlying the activity-induced expression of Emerin, we mined published datasets for Emerin at the transcriptome level. A RiboSeq analysis detected a VE-dependent increase in neuronal *Emd* (the gene encoding Emerin) [12], however, scRNAseq analysis did not identify increased *Emd* in neurons within approximately the same timeframe following VE [51]. These data suggest a potential transcription-independent mechanism might regulate Emerin’s activity-dependent induction. To examine this hypothesis, we first measured *Emd* mRNA level in neuronal cultures at the same timepoints that we had measured Emerin protein in Figure 2E (Figure 3A). *Emd* mRNA remained unchanged 1h and 3h after bicuculline stimulation, when protein had increased significantly as indicated in Figure 2D-E. It was only at 6h after stimulation that we observed a bicuculline-induced increase in *Emd* mRNA (Figure 3B). As positive controls, a panel of immediate early genes, including *cFos* (Figure 3C), *Npas4* (Figure 3D), *Egr1* (Figure S4A) and *Arc* (Figure S4B), showed induction by 1-3h. The temporal dynamics of *Emd* mRNA and Emerin protein levels suggest that the acute induction of Emerin is likely driven by the translation of existing *Emd* mRNA and that activity-dependent *Emd* transcription starts at later timepoints.

**Figure 3.**
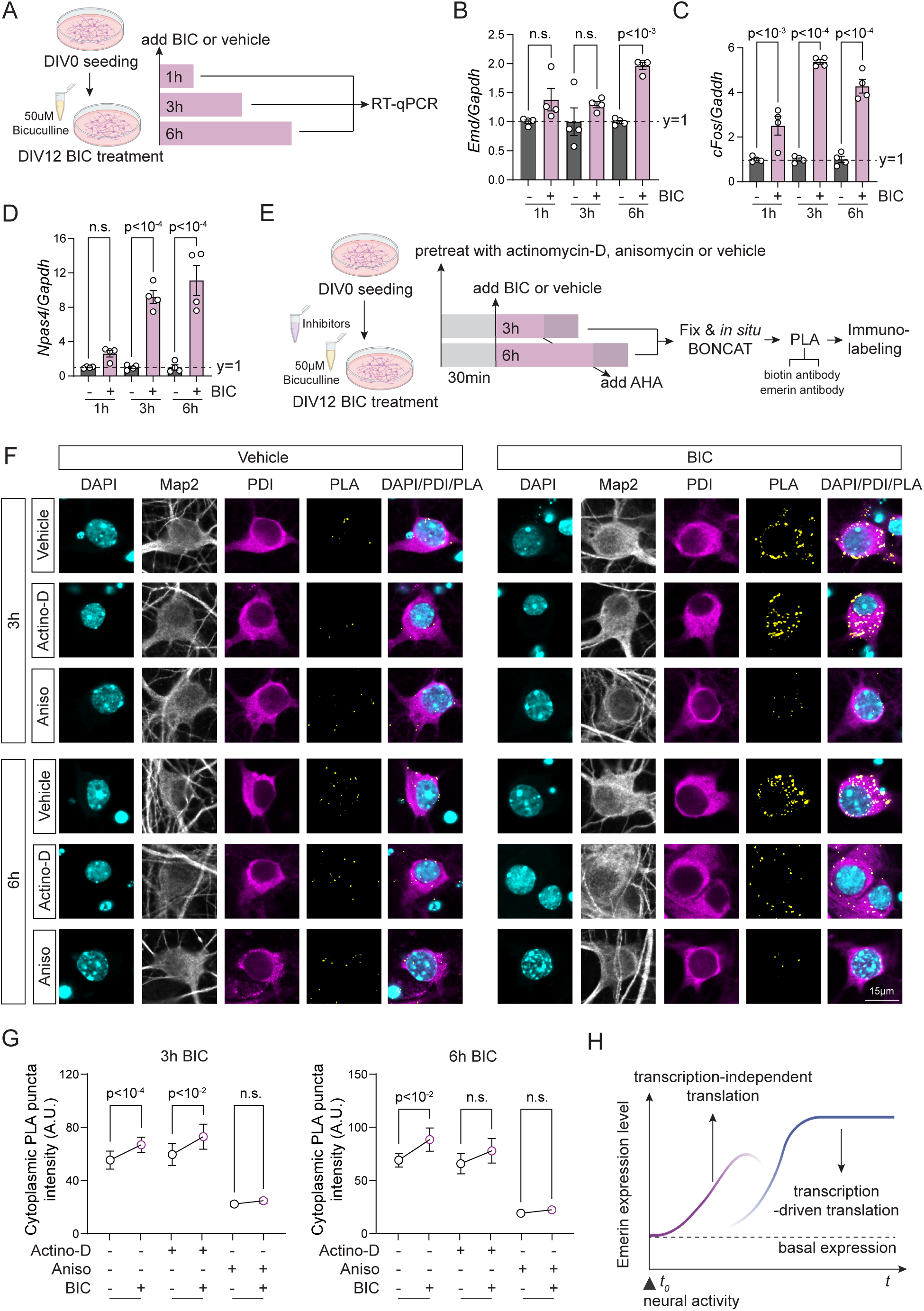
Acute activity-induced Emerin expression is driven directly by translation. (A-D) Timecourse of activity-induced *Emd* mRNA expression. (A) Experimental workflow to assess BIC-induced *Emd* mRNA at 1h, 3h and 6h using RT-qPCR. (B-D) mRNA levels of *Emd* (B) *cFos*(C) and *Npas4* (D) normalized to *Gapdh*. BIC groups are normalized to the corresponding vehicle controls. n=4 individual wells from 2 independent culture preparations. One-way ANOVA test with Sidak multiple comparison (compare BIC to the corresponding vehicle). Bars and lines indicate mean and SEM. n.s., not significant. (E-G) Actinomycin and anisomycin differentially affect the timecourse of activity-induced Emerin synthesis. (E) Experimental workflow to assess effects of actinomycin-D or anisomycin on BIC-induced newly-synthesized Emerin at 3h and 6h using in situ BONCAT-PLA. (F) Representative images of nascent Emerin PLA puncta after BIC/vehicle treatment at each timepoint following actinomycin-D or anisomycin pretreatment. (G) Quantification of cytoplasmic Emerin PLA intensity. BIC groups are normalized to the corresponding vehicle controls. n=1000-1500 individual neurons from 6 wells of 2 independent culture preparations. Kruskal-Wallis test with Dunn’s multiple comparison (compare BIC to the corresponding vehicle). Dots and lines indicate median and 95% CI. n.s., not significant. (H) Schematic summary showing rapid activity-induced Emerin protein synthesis is independent of Emd transcription.

To further unravel the transcriptional and translational controls of activity-induced Emerin synthesis, we pre-treated neuronal cultures with the transcription inhibitor, actinomycin-D, or the translation inhibitor, anisomycin, before bicuculline stimulation. We then measured nuclear and cytoplasmic nascent Emerin puncta using *in situ* BONCAT-PLA (Figure 3E). Bicuculline alone increased nascent Emerin puncta in cytoplasmic compartments but not in the nucleus over 3h and 6h, consistent with the data in Figure 2H. Blocking transcription with actinomycin-D did not affect the bicuculline-induced cytoplasmic increase in Emerin synthesis after 3h but blocked this activity-induced increase after 6h, consistent with our finding that bicuculline-induced *Emd* mRNA increases at 6h (Figure 3B). By contrast, blocking translation with anisomycin significantly reduced Emerin synthesis at both 3h and 6h timepoints (Figure 3F-G, S4C). These data suggest that activity rapidly induces the translation of existing *Emd* mRNA and that transcription-dependent translation drives Emerin synthesis at a later timepoint, corroborating the rapid bicuculline-induced increase in Emerin protein (Figure 2E) and the delayed increase in *Emd* mRNA (Figure 3B). We note that the level of basal Emerin synthesis dropped dramatically with anisomycin treatment, but not with actinomycin-D, suggesting that basal Emerin protein synthesis might rely on transcription-independent translation as well. Together, our data suggest that activity-induced Emerin protein synthesis is rapidly driven by translation, independent of transcription, suggesting that a pool of Emerin mRNA is available for translation in response to an activity-induced signal (Figure 3H).

### Emerin levels bidirectionally affect protein synthesis

Next, we explored the functions of activity-induced Emerin. Emerin and Eif2ak3, a kinase that inhibits translation, interact in the ER [52]. Moreover, the loss of Emerin or the expression of disease-associated Emerin mutants hyperactivates multiple MAPK signaling pathways, including p38-MAPK, ERK1/2 and JNK [53]. These data suggest activity-induced Emerin may play a role in regulating protein translation. To test this hypothesis, we designed, packaged and validated AAV constructs that (1) knockdown Emerin through a cocktail of shRNAs that targets the 3’-UTR, coding sequence and 5’-UTR (a mixture of AAV-U6-shEmd1/2/3, referred to as ‘KD’ hereafter), (2) overexpress Emerin-mRuby fusion protein in a cre-dependent manner in neurons (AAV-hSyn-FLEx-Emerin-mRuby, referred to as ‘OE’ hereafter), (3) express cre recombinase in excitatory neurons (AAV-CaMKIIα-cre) or inhibitory neurons (AAV-vGat-cre) and (4) express the corresponding OE/KD controls (AAV-hSyn-FLEx-mRuby and AAV-U6-shScramble) (Figure S5A-D). Interestingly, when testing our Emerin-OE AAVs, we consistently observed lowered expression of co-transduced AAV-EF1α-FLEx-EGFP, while total protein amount and cultured neuron density remained unaffected (Figure S5E-J).

To test whether Emerin affects protein synthesis, we labeled newly-synthesized proteins in cultures with 1 h AHA treatment and first assessed the amount of labeled nascent protein with BONCAT and western blot (Figure 4A). Emerin KD increased labeled proteins ∼2-fold compared to controls, while Emerin OE in either excitatory or inhibitory neurons decreased biotin-labeled nascent proteins by 50% (Figure 4B-C). These data suggest that low levels of Emerin increase protein synthesis and elevated Emerin levels decrease protein synthesis. Furthermore, Emerin KD partially rescued the Emerin upregulation-induced decrease in protein synthesis, indicating that Emerin regulates protein synthesis bidirectionally (Figure S5K-M). Next, we assessed neuron-specific changes in nascent proteins with single-cell resolution using fluorescent noncanonical amino acid tagging (FUNCAT, Figure S1B bottom), together with visualization of Emerin expression levels by either mRuby intensity in Emerin OE conditions or endogenous Emerin immunolabeling in Emerin KD conditions (Figure 4A). Emerin KD increased the FUNCAT signal while Emerin OE in either excitatory or inhibitory neurons decreased the FUNCAT signal, similar to the BONCAT results, indicating that Emerin levels bidirectionally affect neuronal protein synthesis (Figure 4D-G). Comparing neighboring neurons, FUNCAT signal intensity inversely correlated with Emerin levels, suggesting a cell-autonomous and dose-dependent effect of Emerin expression on neuronal protein synthesis.

**Figure 4.**
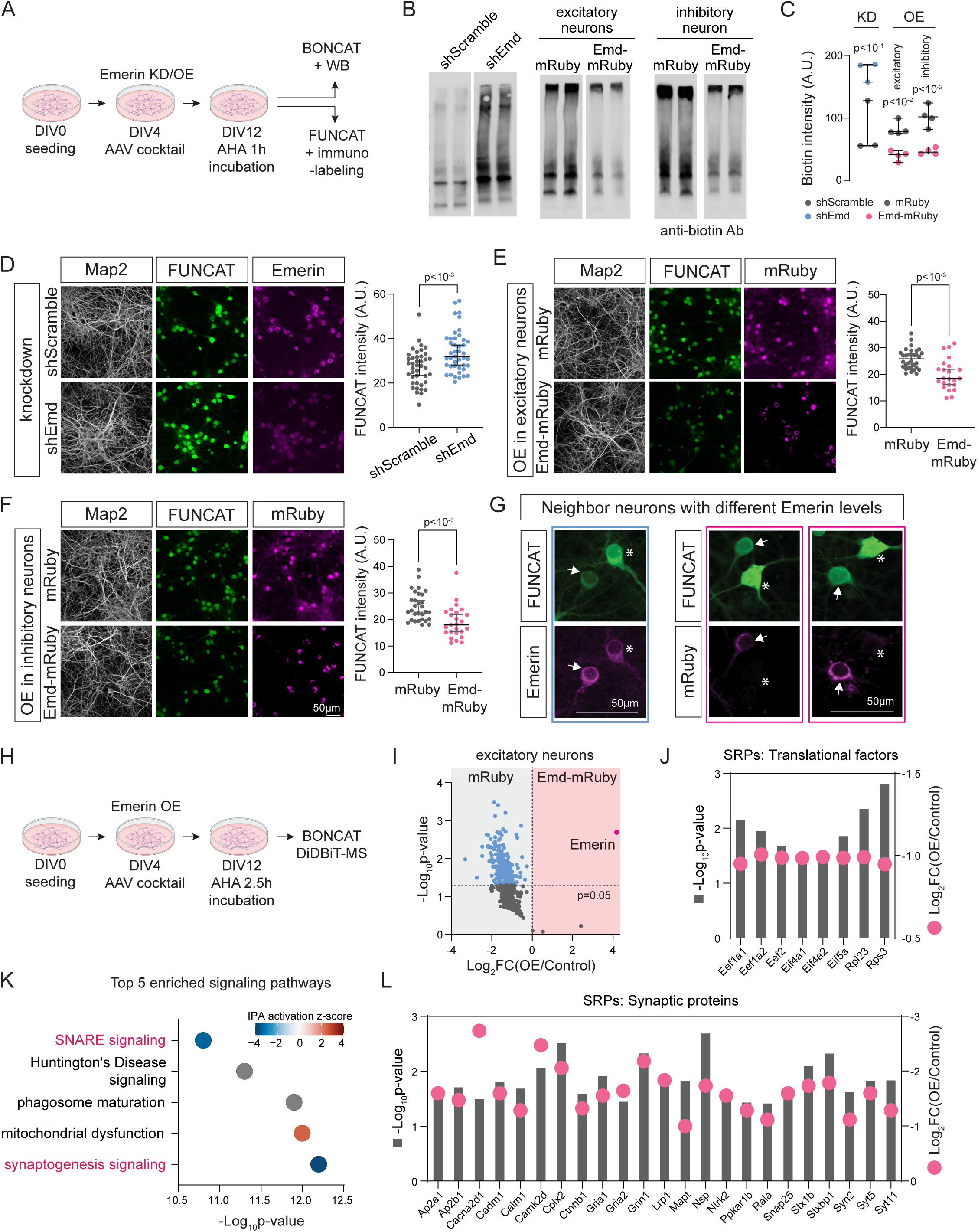
Emerin bidirectionally regulates protein synthesis including translation factors and synaptic proteins. (A-C) BONCAT-western blot (BONCAT-WB) analysis of newly-synthesized proteins with Emerin KD and OE. (A) Experimental workflow for BONCAT and FUNCAT. DIV12 neuronal cultures were treated with AHA for 1h, then subjected to BONCAT-WB or FUNCAT-immunolabeling. (B) BONCAT-WB biotin signal with Emerin KD (shEmd vs control shScramble) (Left), or OE (Emd-mRuby vs control mRuby) in excitatory (middle) and inhibitory neurons (right). (C) Quantification of biotin intensity (normalized to ponceau). n=3-4 individual wells from 2 independent culture preparations. Welch’s t-test. Lines indicate median and 95% CI. (D-G) FUNCAT analysis of newly-synthesized proteins with Emerin KD and OE. Representative FUNCAT images (Left) and statistical analysis with Emerin KD (D), Emerin OE in excitatory (E) and inhibitory (F) neurons. (G) Enlarged images showing neighboring neurons in experimental groups with different Emerin expression levels. Arrow, neurons with high Emerin level. Asterisk, neurons with low Emerin level. (G) Quantification of FUNCAT signal. n=24-33 individual neurons analyzed from 4 wells. Welch’s t-test. Lines indicate median and 95% CI. (H-L) Newly-synthesized proteome with Emerin OE. (H) BONCAT-DiDBiT-MS/MS experimental workflow. (I) Volcano plot of all biotinylated proteins identified. n=4 biological replicates. (J) Emerin OE-regulated proteins that are translational factors. (K) Top 5 significantly enriched IPA pathways. Magenta, synapse-related pathways. (L) Emerin OE-regulated proteins that are synaptic proteins.

To determine the targets of Emerin’s translational control, we employed our proteomics pipeline to identify the nascent proteome labeled with 2.5h exposure to AHA in Emerin OE neurons (Figure 4H). Proteomic analysis identified a total of 634 newly-synthesized proteins, 206 of which were significantly down-regulated by Emerin OE (Figure 4I, Table S4). A group of 8 translational regulators were downregulated, including eIF4a, eIF5a, eEF1a and eEF2, consistent with the protein synthesis inhibition phenotype seen with increased Emerin expression (Figure 4J). IPA analysis identified synaptogenesis as the top pathway regulated by increased Emerin expression and predicted that increased Emerin would negatively regulate this pathway based on the decreased synthesis of 23 synaptic proteins (Figure 4K-L). Together, our data suggest that activity-induced Emerin upregulation might lead to a downstream decrease in neuronal protein synthesis and predict that altering Emerin expression may lead to a synaptic phenotype, which we examine in our next experiments.

### Emerin regulates synapse density and spine morphology

We first evaluated the effect of Emerin KD and OE on synaptic density by immunolabeling Synaptophysin and PSD95 to identify pre-and postsynaptic elements, respectively (Figure 5A, 5B-left). Presynaptic and postsynaptic puncta were quantified, and synapses were identified based on their colocalization. Synaptic density increased with Emerin KD and decreased with Emerin OE in excitatory or inhibitory neurons, consistent with the IPA prediction from the Emerin OE nascent proteome (Figure 5D-E). When examining the synaptic components individually, we found that the density of postsynaptic marker PSD95 was bidirectionally affected by Emerin levels, whereas the density of presynaptic marker Synaptophysin remained unchanged. To further dissect the synaptic phenotypes, we immunolabeled markers specific to excitatory synapses (vGlut1 and Homer1) and inhibitory synapses (vGat and Gephyrin) (Figure 5B, right). Both excitatory and inhibitory presynaptic markers (vGlut1 and vGat) were unaffected by Emerin manipulations (Figure S6A-C). By contrast, Homer1 puncta density increased with Emerin KD and decreased with Emerin OE (Figure 5F-H, S6D), following the same trend as PSD95. It’s interesting to note that Gephyrin puncta density was unaffected by Emerin KD but increased with Emerin OE, suggesting that increased Emerin expression may have opposite effects on excitatory and inhibitory postsynaptic sites.

**Figure 5.**
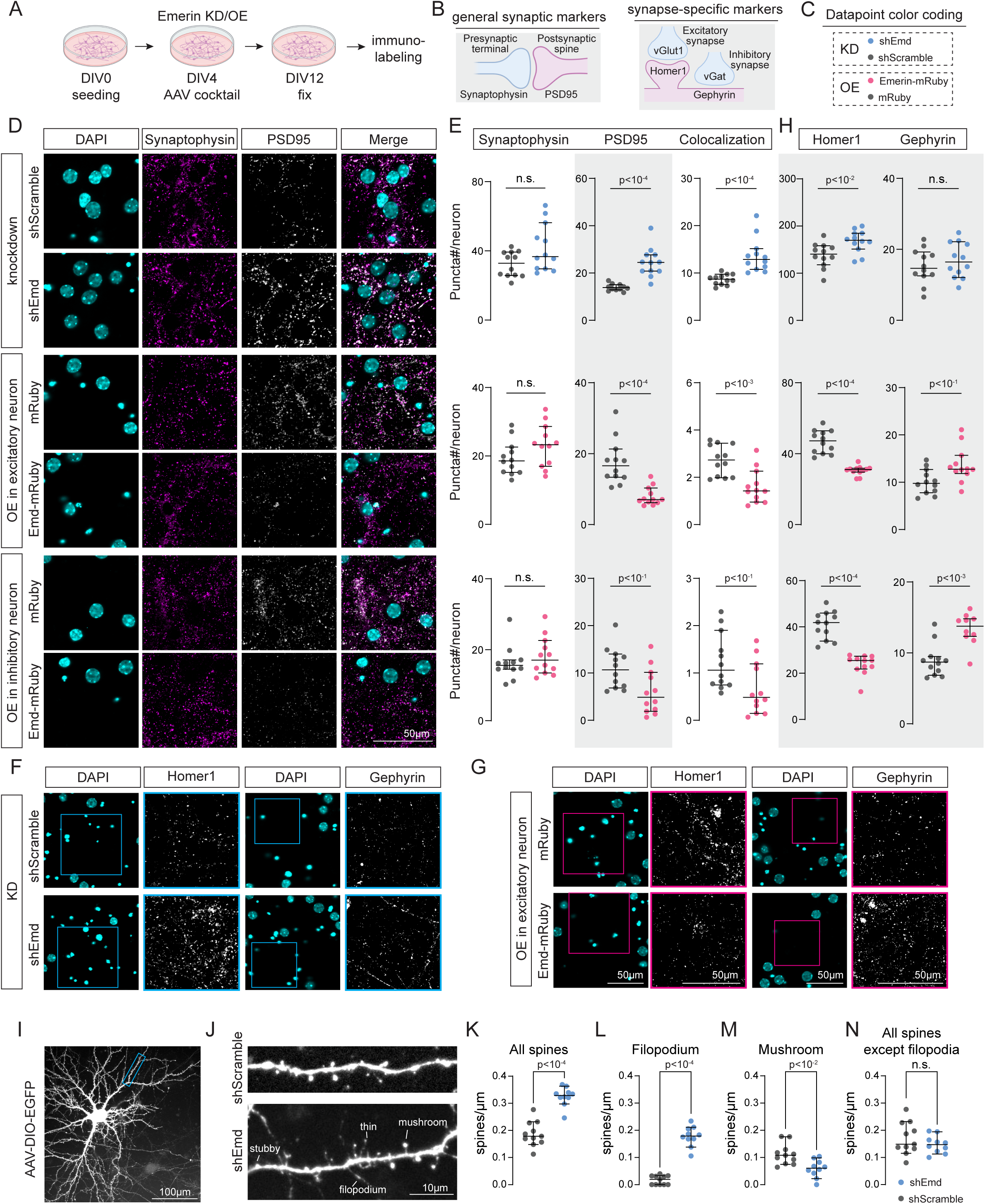
Emerin regulates synapse density and spine morphology. (A-H) Synaptic density analysis with Emerin KD and OE. (A, B) Experimental workflow (A). DIV12 neuronal cultures were immunolabeled with synaptic markers (B). (C) Color-code for data in E, H. (D) Representative images of Synaptophysin and PSD95 immunolabeling with Emerin KD (top 2 rows), Emerin OE in excitatory (middle 2 rows) and inhibitory neurons (bottom 2 rows). (E) Quantification of Synaptophysin and PSD95 puncta and their colocalization with Emerin KD (top row), Emerin OE in excitatory (middle row) and inhibitory neurons (bottom row). (F,G) Representative images of Homer1 and Gephyrin with Emerin KD (F) and Emerin OE in excitatory neurons (G). Boxed regions in DAPI images are shown enlarged for Homer1 and Gephyrin. (H) Quantification of Homer1 and Gephyrin puncta with Emerin KD (top row), Emerin OE in excitatory (middle row) and inhibitory neurons (bottom row). Quantification on postsynaptic components in E,H were highlighted in grey. Puncta counts are normalized to neuron numbers, based on DAPI. n=12 individual wells from 4 independent culture preparations, Mann-Whitney test. Lines indicate median and 95% CI. n.s., not significant. (I-N) Spine morphology analysis with Emerin KD. (I) Representative image of a sparsely-labeled neuron expressing EGFP. Blue rectangle, a representative secondary dendrite selected for analysis. (J) Representative ∼45µm dendritic stretches from Emerin KD and sham KD neurons. Examples of the four spine types quantified are labeled. Quantification of spine counts of all types (K), filopodia (L), mushroom (M) and all types excluding filopodia (N). n=10-11 dendritic stretches, each from one individual neuron. 3 independent culture preparations, Mann-Whitney test. Lines indicate median and 95% CI. n.s., not significant.

To dissect the Emerin KD-induced increase in excitatory synaptic contacts, we expressed EGFP in neurons to visualize their spine morphology (Figure 5I). We quantified the density of 4 distinct spine types on neuronal dendrites: thin, stubby, mushroom and filopodium (Figure 5J). Total spine density significantly increased with Emerin KD, recapitulating the increase in PSD95 and Homer1 densities (Figure 5K). Surprisingly, the analysis of individual spine types showed that filopodial spines were the only type that significantly increased in density with Emerin KD, whereas mushroom spine density decreased (Figure 5L-M, S6E-F). Thin and stubby spines did not change significantly. Total spine quantification, excluding filopodia, showed no difference between Emerin KD and control, indicating that the knockdown-induced increase in spine density was primarily due to increased filopodia (Figure 5N). The increase in filopodial spine density and the decrease in mushroom spine density suggest a net shift of the spine population from strong mushroom spines to weak filopodial spines with Emerin KD and further suggest that activity-induced Emerin upregulation might reduce immature filopodial spines and promote mature mushroom spines. Moreover, the increase in Gephyrin immunolabeling with Emerin OE suggests that activity-induced Emerin upregulation might also increase inhibitory synaptic contacts (Figure 5G, S6D). Together, our data suggest that increased Emerin expression promotes neuronal connectivity by regulating the density and morphology of postsynaptic elements, which could be manifested as enhanced neural network activity. We test this hypothesis in our next experiments.

### Emerin is required for normal visual responses in mice

The data described above suggest a role for Emerin in a feed-forward effect of activity on network function. To examine this hypothesis, we first measured spontaneous activity in neuronal cultures, by expressing the calcium indicator GCaMP7s in excitatory or inhibitory neurons in a cre-dependent manner. We recorded calcium transients over 100s epochs. Emerin KD decreased the number of spontaneous calcium transients during each epoch in both excitatory and inhibitory neurons (Figure S7A-B), whereas Emerin OE increased spontaneous calcium transients in both cell types (Figure S7C-D). These data suggest that Emerin levels bidirectionally alter spontaneous network activity in vitro, which prompted us to hypothesize that Emerin might regulate network properties in response to in vivo sensory experience as well.

To test this, we unilaterally knocked-down Emerin in mouse primary visual cortex (Figure S8A-B) and bilaterally measured visual stimulation-evoked local field potentials (VEPs). The unilateral AAV-shEmd injections were alternated between the left and right hemispheres to exclude any hemisphere-specific effect, with AAV-shScramble injected into the contralateral cortex as the control (Figure 6A-B). Mice were dark-adapted for 1 hour before recording to prime retinal cells to visual stimulation [54], anesthetized and provided with visual stimulation with an LED stroboscope at 1Hz for 250 iterations (Figure 6C-D). Emerin KD decreased VEP trough-to-peak amplitude by 36% compared to the AAV-shScramble control from the contralateral cortex, indicating that Emerin KD reduced visual responses (Figure 6E-F). Data compared between the left and right hemispheres were not significantly different (Figure 6F). Interestingly, we also observed a significant increase in VEP latency with Emerin KD (Figure 6G). This small increase in response kinetics (Δ=1.2ms) might result from synaptic remodeling, such as an increase in weaker filopodial spines within the visual cortex as a result of Emerin KD.

**Figure 6.**
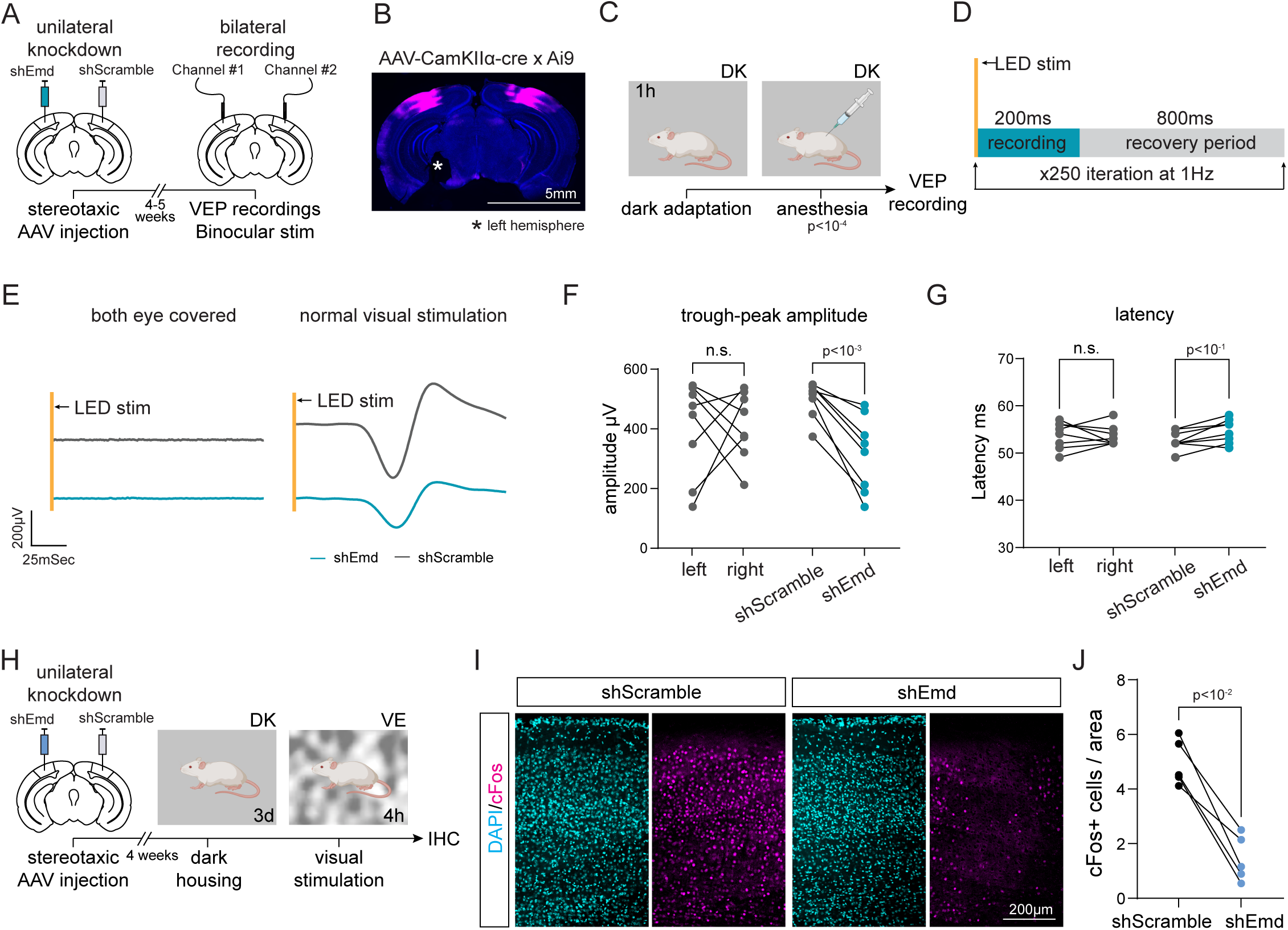
Emerin is required for normal visual response in mice. (A-G) Visually-evoked potential (VEP) recordings with Emerin KD. (A) Experimental workflow of unilateral Emerin knockdown and bilateral VEPs recording. (B) Representative coronal brain image showing AAV targeting and coverage. Left hemispheres are marked during sectioning for *post-hoc* data interpretation. (C) Workflow for pre-recording preparation. (D) Visual stimulation and data acquisition protocol. (E) Representative VEP traces when both eyes were covered (left) and with normal visual stimulation (right). Recordings from KD (blue) and control (grey) hemispheres are aligned to the stimulus. (F) Quantification of trough-peak amplitude comparing left vs right hemispheres (left) and comparing shEmd vs shScramble treatments (right). (G) Quantification of VEP latency comparing left vs right hemispheres (left) and comparing shEmd vs shScramble treatments (right). n=8 mice, paired t-test. n.s., not significant. (H-J) Visual experience-induced cFos expression with Emerin KD. (H) Experimental workflow of unilateral Emerin knockdown and visual experience paradigm. (I) Representative image of VE-induced cFos immunolabeling in primary visual cortex. (J) Quantification of cFos+ neurons comparing shEmd and shScramble treatments. n=5 mice, paired t-test.

We further tested whether Emerin KD affected visual stimulation-induced cFos expression as an independent indicator of Emerin’s role in visual cortical responses to sensory input. We adopted a similar strategy of unilateral Emerin KD as described above for VEPs recordings. Mice were subjected to 3-day dark-housing followed by 4h of visual stimulation (Figure 6H). Emerin KD significantly reduced the number of cFos-positive neurons following visual stimulation compared to the shScramble-treated cortex (Figure 6I-J). Together, these results indicate that Emerin is essential for normal visual stimulation-induced responses in mice.

### Emerin is required for visual information processing in mice

Finally, we tested if Emerin function is required for visual information processing. We used the visual cliff test to evaluate depth perception in mice, which involves neuronal integration of binocular information in the primary visual cortex [55]. The visual cliff arena provides visual cues at different depths below a transparent platform, segregating the perception of ‘safe’ and ‘unsafe’ zones in the arena (Figure 7A). Mice were injected bilaterally with AAV-shEmd or AAV-shScramble in the visual cortex and were subjected to two test sessions in the visual cliff arena, one at day 5 and the other at day 30 after surgery (Figure 7B). Location tracking of mice assayed at day 30 showed that the control mice avoided the unsafe zone, whereas the Emerin KD mice displayed no avoidance of the unsafe zone and moved between the safe and unsafe zones more often than the controls (Figure 7C). The color-coded location tracking data of the mice indicated that the Emerin KD mice spent significantly less time in the safe zone than control mice (Figure 7D). Quantitatively, these Emerin KD mice spent 25% and 75% of the time, on average, in the safe and unsafe zones respectively, reflecting the 25-75 area division of the two zones (Figure 7E). This indicates that the Emerin KD mice failed to detect or respond to the depth cues. Furthermore, Emerin KD resulted in increased distance traveled in the unsafe zone and increased average traveling speed in the safe zone, suggesting disrupted visual cue-guided behaviors by Emerin KD (Figure 7F-G). Comparing the time spent in the safe zone at day 5 and day 30 showed a decline in performance of the KD mice over time, while the behavior of the control mice remained constant (Figure 7H). Taken together, these experiments indicate that Emerin is required for visual information processing underlying depth perception in mice.

**Figure 7.**
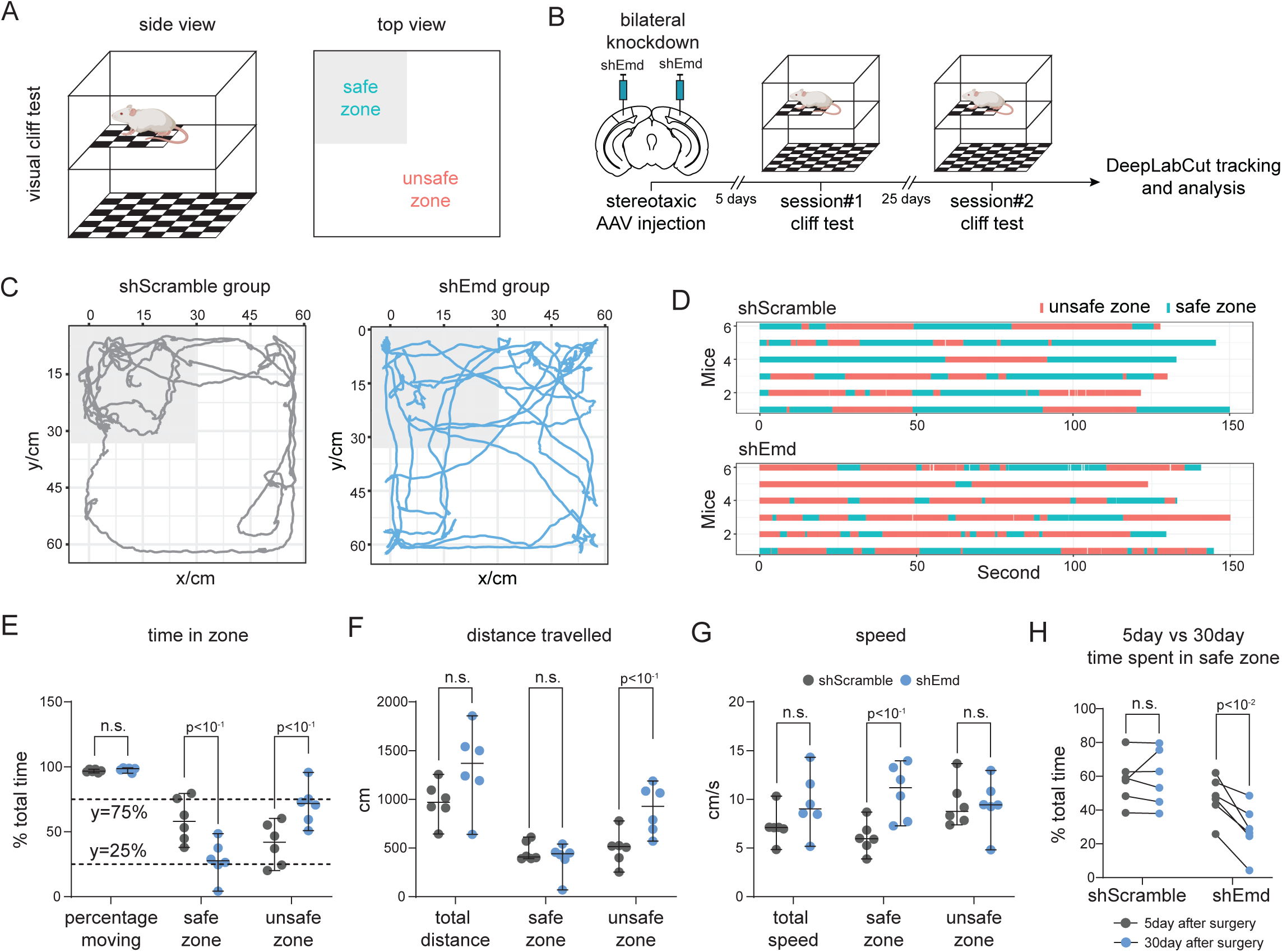
Emerin is required for visual depth perception in mice. (A) Side view schematic of the visual cliff arena (Left) and top view showing the 25/75 distribution of safe and unsafe zones (Right). (B) Experimental workflow of bilateral Emerin knockdown and visual cliff test sessions 5 days and 30 days after AAV injection. (C) Representative movement tracking of mice within the visual cliff arena using DeepLabCut. Safe zone is highlighted in grey. (D) Raster plots of time spent in safe (green) and unsafe (orange) zones of the shEmd and shScramble groups. (E-G) Quantification of behavioral parameters from the day 30 session. (E) Percent total time spent moving and relative time spent in the safe and unsafe zones. (F) Quantification of distance travelled in the entire arena, safe zone, and unsafe zone. (G) Quantification of average moving speed in the entire arena, safe zone, and unsafe zone. n=6 mice, Welch’s t-test between shEmd and shScramble. Lines indicate median and 95% CI. n.s., not significant. (H) Quantification of time spent in the safe zone comparing day 5 and day 30 after AAV injection. n=6 mice, paired t-test. n.s., not significant.

## Discussion

Alterations in the newly-synthesized proteome can be influenced not only by transcriptomic changes [21–24] but also by multiple layers of post-transcriptional regulation of protein synthesis [25–29], trafficking [30, 31] and degradation [32, 33]. Consequently, the nascent proteome in the brain of intact animals is highly dynamic over time after exposure to external stimuli. A direct proteomic evaluation of the visual experience-induced nascent proteome within an early and defined time window after stimulation could lead to new discoveries of protein effectors of neuronal plasticity. Here, we applied a neuronal cell type-specific BONCAT-DiDBiT-MS/MS proteomics pipeline and enhanced its temporal resolution by increasing short-term labeling efficiency. This optimized pipeline allowed us to detect the cell type-specific expression of 2883 proteins with high confidence, within 10h of the onset of visual experience. This temporal scale provides the opportunity to identify rapid in vivo changes in the nascent proteome, including candidate plasticity proteins which may function as immediate early effector proteins. Visual experience-dependent plasticity has distinct features during the developmental critical period and in the adult and in different neuronal populations [4–6, 56], suggesting diverse molecular programs mediate age-and cell type-dependent plasticity. We found distinct visual experience-induced proteomic signatures in EX28, IN28, EX56 and IN56, with less than 30% CPP overlap and moderate CPP correlations across samples, providing a rich resource of candidates for further functional investigation. Moreover, a bioinformatic search of shared mechanisms across all conditions revealed broad recruitment of proteostasis pathways. Building on the recognized significance of proteostasis in maintaining neuronal and circuit function, our data suggest that this regulation might have an activity-dependent component in a broad range of cellular contexts.

We took an effect size-based, unbiased approach to select Emerin, the top CPP, for functional evaluation. Both the proteomic screen and independent validation *in vitro* and *in vivo* demonstrated Emerin’s strong activity-induced synthesis, consistent with detecting Emerin in our previous study of seizure-induced changes in the nascent proteome [39]. Interestingly, acute activity-dependent Emerin synthesis is insensitive to inhibiting transcription, suggesting that this rapid Emerin synthesis relies on translation from an existing *Emd* mRNA pool and is independent of newly-transcribed *Emd* or other genes. These data suggest that Emerin is an immediate early effector protein, analogous to the immediate early effector genes such as CPG15, CPG2 [57], Homer1 [58] and Arc [59]. This independence between Emerin’s activity-induced translation and transcription echoes a growing body of evidence indicating that the complex regulation of proteome content might be achieved through independently targeting (post-)transcriptional and (post-)translational mechanisms [60–62].

Emerin is an integral membrane protein translated in the cytosol and post-translationally inserted into the ER membrane [41, 42]. Emd mRNA is a target of many RNA-binding proteins (RBP), including RMRP [63], Pumilio [64], MOV10 and NXF1 [65]. Furthermore, Emd mRNA is predicted to have binding sites for a large pool of RBPs by both sequence-(RBPmap) and structure-based (POSTAR3) algorithms (data not shown). RBPs are sensitive to neuronal activity [66], which could be a regulatory mechanism for activity-dependent transcription-independent Emerin synthesis.

The majority of Emerin studies focused on muscle, where it has diverse functions relating to its subcellular localization [45, 67–69]. Muscle Emerin is predominantly located in the inner nuclear envelope based on its high affinity association with Lamin A [41–43], where it regulates signaling, gene expression, and nuclear and chromatin structure [45]. By contrast, we find that in neurons, which lack Lamin A [48], both basal and activity-induced newly-synthesized Emerin are concentrated in the extranuclear somatic compartment, likely in the ER considering Emerin’s biogenesis and colocalization with the ER marker PDI.

The ER plays an essential role in proteostasis through the regulation of protein synthesis, folding and trafficking [70–72]. ER function in the synthesis of transmembrane proteins and their trafficking to the plasma membrane is particularly salient in neurons, as exemplified by its role trafficking glutamate receptors following induction of long-term synaptic plasticity [70, 71]. Upregulating Emerin, which mimics activity-induced increased Emerin expression, robustly decreased protein synthesis, whereas downregulating Emerin, which could mimic ER stress induced degradation of Emerin [73], increased protein synthesis in a cell autonomous manner. Given the enrichment of both activity-induced neuronal Emerin and translation associated machinery in the ER, this bidirectional regulation of protein synthesis may be regulated by ER-associated Emerin. An interactome analysis of Emerin indicates that it associates with numerous ER proteins [50], including eIF2ak3/PERK, an ER kinase and translation inhibitor. Enrichr bioinformatic analysis linked Emerin and its co-expressed network to translation, ribosomes and mRNA binding (Table S5). In Emerin knockout myocytes, MAPK signaling pathways are activated [74–77], consistent with the observed increase in translation we see with Emerin knockdown. Our proteomic analysis under Emerin OE conditions indicated that >60% of the down-regulated proteins are membrane proteins. This evidence collectively suggests that Emerin might influence protein synthesis in the ER by affecting MAPK pathways, translation factors and ribosome complexes, though pinpointing the exact mechanisms requires more in-depth analysis [73, 78–82]. Alternately, the feed forward Emerin-dependent regulation of protein synthesis could be attributed to Emerin’s function in the inner nuclear envelope, where studies in myocytes show that molecular partners such as HDAC3 are recruited by Emerin and repress gene expression [83]. Our MS/MS proteomic analysis of the proteins that were down-regulated in response to increased Emerin identified synaptic proteins and translational regulators. This indicates that the activity-dependent increase in Emerin orchestrates a downstream wave of activity-induced proteome dynamics through negative regulation of protein synthesis, providing a potential brake on activity-induced translation and a temporal switch on proteostasis regulation in response to sensory input.

Many regulators of activity-dependent plasticity mediate the addition or elimination of synapses or affect their structural and functional properties, culminating in circuit-level modifications encoding experience-relevant information [3, 84]. We found that Emerin downregulation resulted in a significant shift in the morphology of the population of spines, increasing filopodial spines and decreasing mushroom spine density. This suggests that activity-induced Emerin may play a role in maintaining efficient neuronal connectivity by coordinating the elimination of weak connections and the maturation of strong, information-encoding connections. Our proteomic data identified a large group of synaptic proteins whose expression was reduced with Emerin upregulation, suggesting Emerin’s regulation of synaptic connectivity might be linked to its activity-dependent regulation of protein synthesis. Though Emerin is concentrated in the neuronal cell body, consistent with somatic enrichment of ER, we observed Emerin immunolabel as well as newly-synthesized Emerin puncta in neuronal processes (data not shown), potentially through active trafficking or local synthesis of Emerin. This raises the interesting possibility that activity-induced Emerin may regulate synaptic properties locally, consistent with reports that Emerin associates with RNA binding proteins, such as Pumilio [64] and MOV10 [65], which are known to also regulate synthesis of synaptic proteins [78, 80].

Activity-dependent synaptic plasticity underlies neural network reorganization and is often reflected in functional circuit properties [7, 8] and animal behavior [15–17]. Considering the multi-organ expression of Emerin throughout the lifespan (Human Protein Atlas [85], https://www.proteinatlas.org/), we leveraged viral knockdown of Emerin to restrict downregulation of Emerin to the adult mouse visual cortex for *in vivo* physiological and behavioral analysis. Downregulating activity-induced Emerin expression reduced the amplitude of VEPs and impaired depth perception-guided cliff avoidance behavior. These data suggest that activity-dependent Emerin synthesis is required to maintain strong visual responses to task-relevant visual input and visual information processing to guide behavior.

Based on Emerin’s sensory experience-driven synthesis and its functions at the molecular, cellular, physiological and behavioral levels, we propose a model in which visual experience induces a rapid increase in Emerin synthesis that strengthens synaptic connectivity, facilitating enhanced network response to visual input and intact visual information processing to guide survival behavior. Emerin serves a feed-forward role in a cascade of regulatory processes by inhibiting protein synthesis, gating activity-dependent neuronal plasticity through coordinating proteostasis.

## Acknowledgements

We thank Cline lab members and alumni for their input and feedback and Dr. Cris Niell for discussions and the visual stimulation code. We are grateful to Dr. Larry Gerace for valuable discussions. We thank the Scripps Research Department of Animal Resources for mouse colony management and technical support and the Dorris Neuroscience Microscopy Core, especially Dr. Kathyrn Spencer for technical support. We thank our colleagues in the Dorris Neuroscience Center, especially the Ye lab, the Maximov lab, and the Lippi lab, for sharing reagents, equipment and knowledge, and Erin McAuliff. and Xuanyu Dong for cloning and genotyping, respectively. Y.X. is supported by the Dorris Scholar award. Erin McAuliff. and Xuanyu Dong. were supported by the UC San Diego Academic Internship Program. This work is supported by the National Institutes of Health Grants EY031597 (to H.T.C.), GM103533 and MH067880 (to J.R.Y.), AG075862 (to H.T.C. and J.R.Y.), and AG 067331 and AG 069206 (to M.P.).

## Author Contributions

H.T.C. and Y.X. conceptualized, designed, and wrote the paper. Y.X. performed all experiments and analyzed the data, in close collaboration with other authors as noted below. R.W. made primary neuronal cultures, performed immunoblotting and histology. D.M., Y.M. and J.D. ran LC-MS/MS. M.S. helped perform VEP recordings. H.T.C., J.R.Y., M.P. managed the projects and supervised the study. All authors reviewed and provided feedback on the manuscript.

## Declaration of Interests

The authors declare no competing financial interests.

---

**Figure S1.**
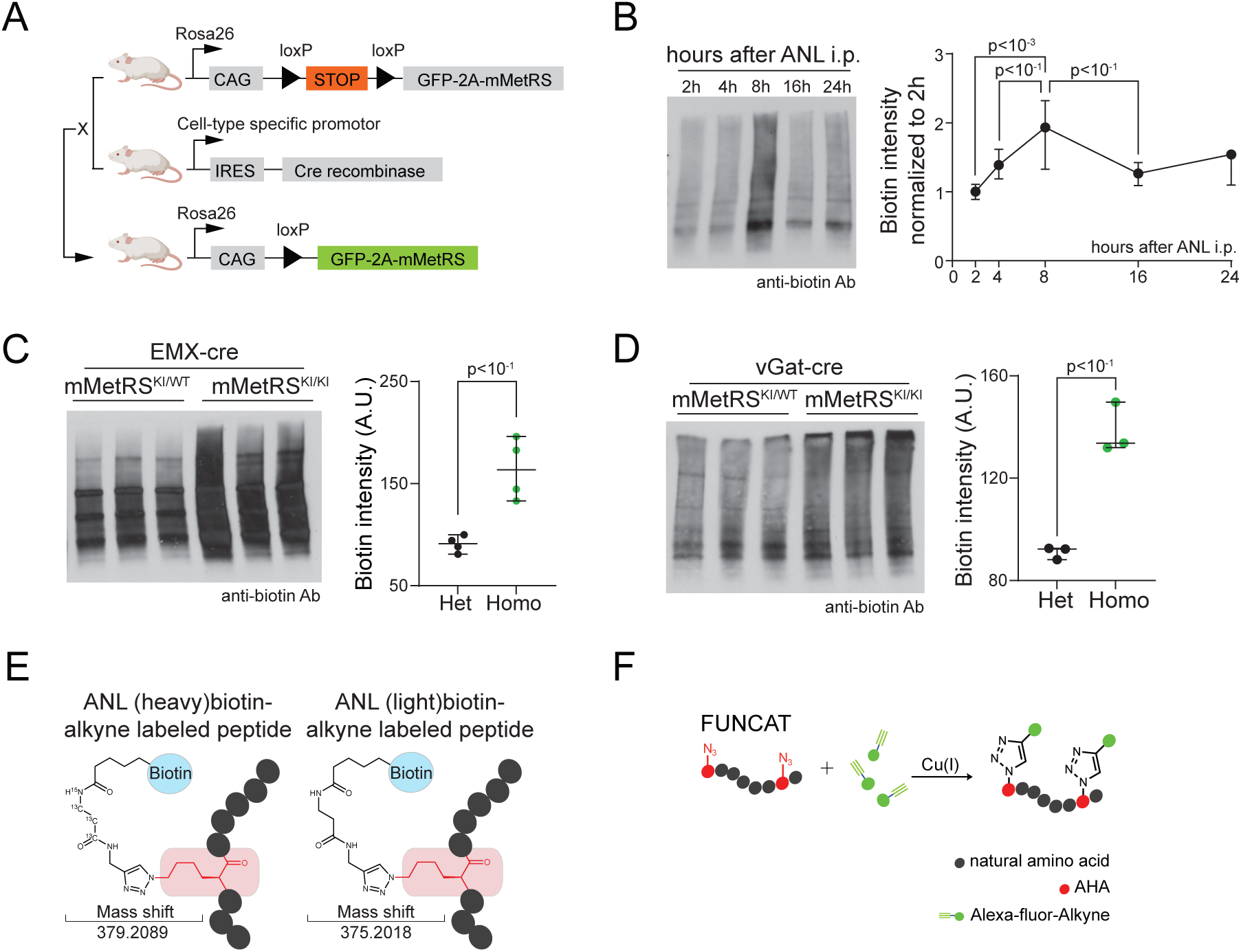
Optimization of cell type-specific nascent proteomics pipeline for studying acute dynamics (related to Figure 1). (A) Breeding scheme to generate mouse models expressing mMetRS in the cell type of interest. F1 offsprings were backcrossed to generate homozygous mMetRS^KI/KI^ (see Methods). (B-D) Optimization of ANL labeling efficiency in mMetRS^KI^ lines. (B) BONCAT-western blot (WB) analysis of ANL incorporation kinetics after one 1.p. injection. Left, representative blot. Right, quantification of biotin intensity. n=3-5, one-way ANOVA with Tukey’s multiple comparison. (C, D) BONCAT-WB analysis comparing ANL labeling efficiency between homozygous mMetRS^KI/KI^ and heterozygous mMetRS^KI/WT^, crossed to EMX^cre^ (C) and vGat^cre^ (D). Tissue from EMX^cre^-mMetRS was collected 16h after i.p. ANL injection. vGat^cre^-mMetRS mice received 2 i.p. injections at a 24h interval and tissue was collected 16h after the second injection. Left, blot showing 3 biological replicates. Right, quantification of biotin intensity. n=3, Welch’s t-test. Lines indicate median and 95% confidence interval (CI). (E) Schematics of proteomics sample duplexing through click chemistry using heavy-and light-isotope modified biotin-alkynes. Differential mass shifts were identified on labeled peptides by MS/MS. (F) Click chemistry schematics for FUNCAT readouts from AHA metabolic labeling experiments.

**Figure S2.**
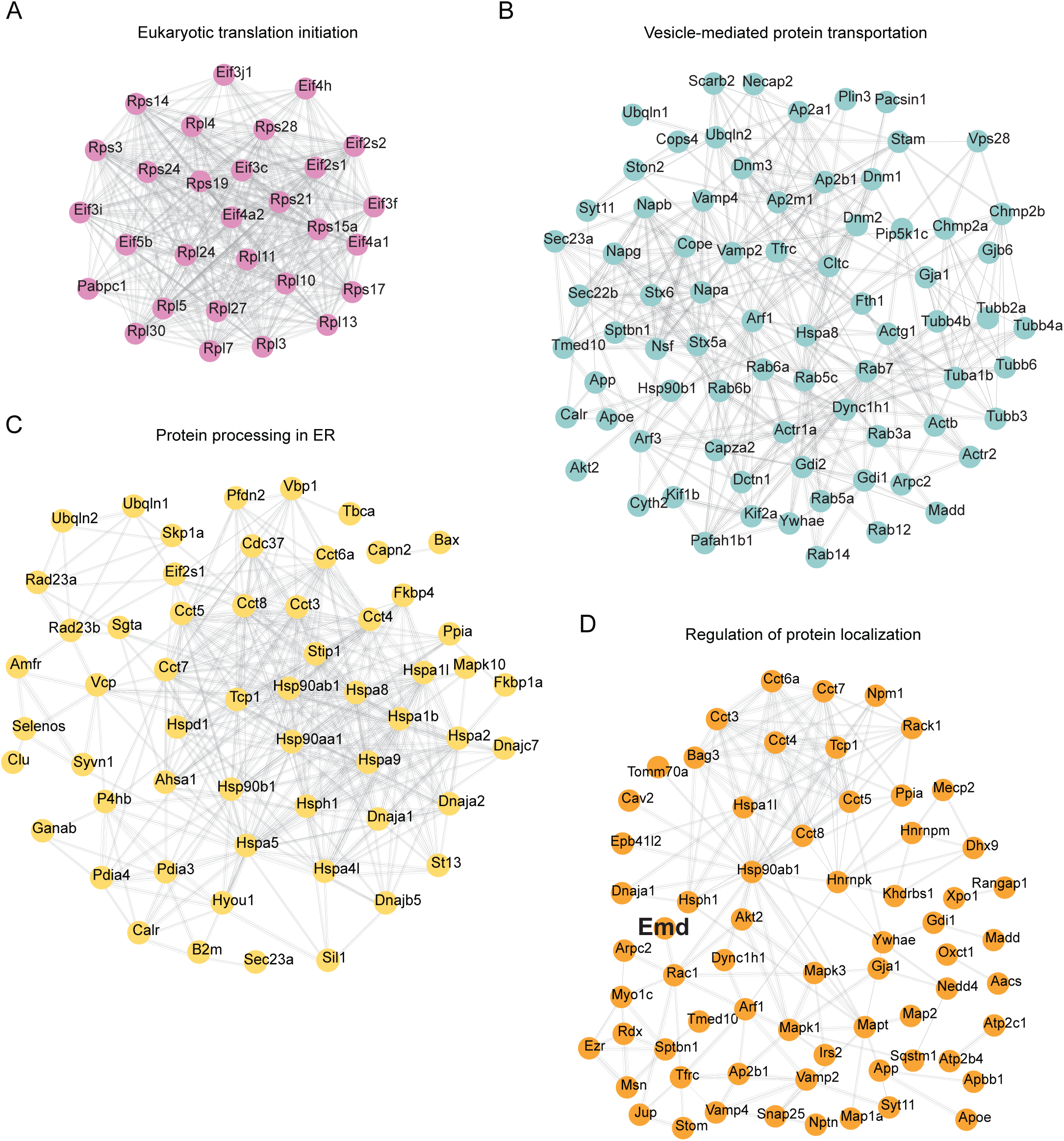
Shared proteostasis-related pathway clusters across ages and cell types identified by Metascape (related to Figure 1). (A-D) Shared Metascape pathways associated with proteostasis regulation. (A) Eukaryotic translation initiation. (B) Vesicle-mediated protein transportation. (C) Protein processing in the ER. (D) Regulation of protein localization. Pathway components are visualized in protein-protein interaction networks by STRING.

**Figure S3.**
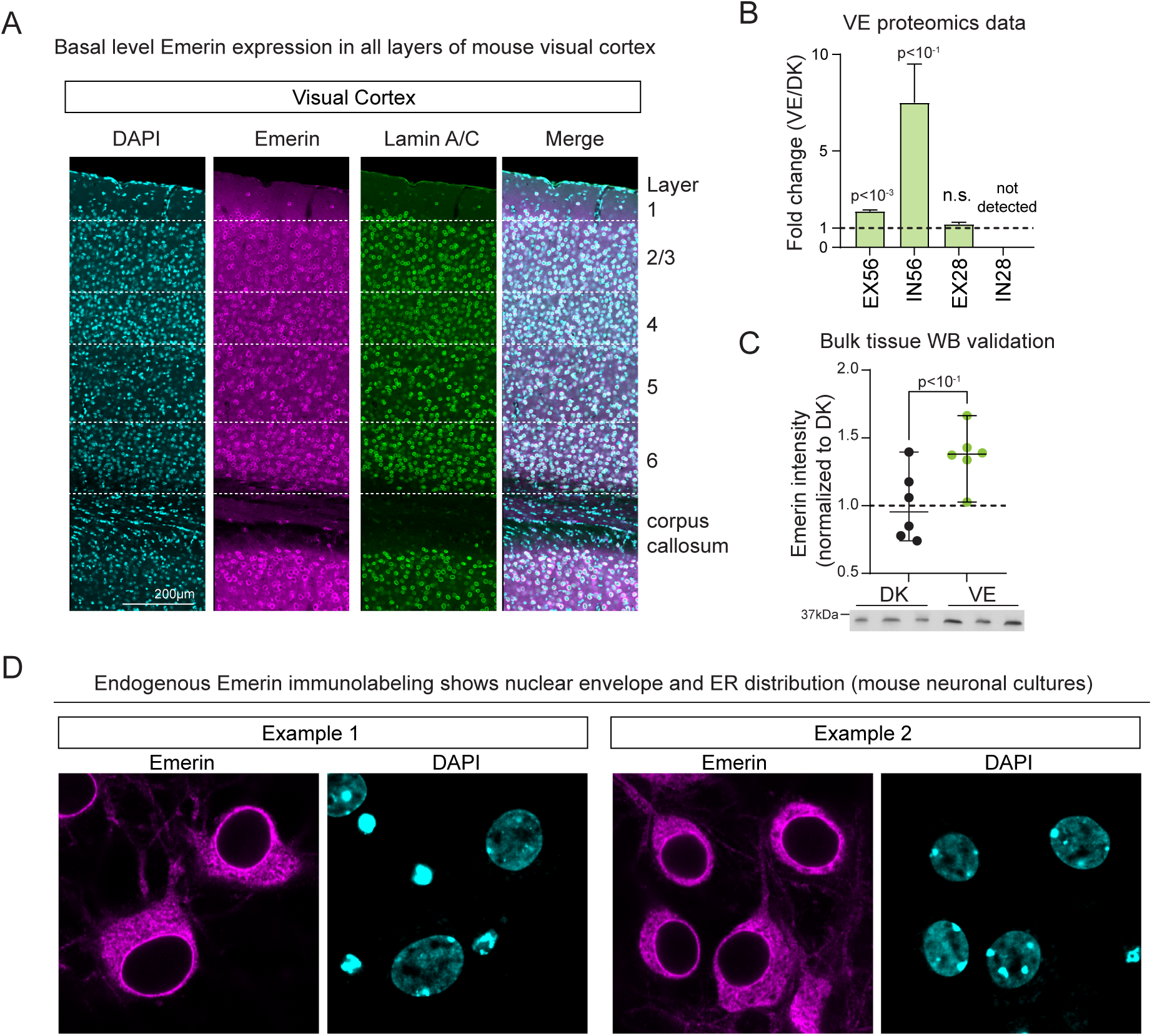
Basal and visual experience-dependent Emerin expression in mouse primary visual cortex (related to Figure 2). (A) Representative image of a coronal section through primary visual cortex immunolabeled for Lamin-A and Emerin and counterstained with DAPI. Emerin immunolabeling is detected in all layers of primary visual cortex in the basal state (12h light/12h dark light cycle). White dotted lines indicate layer boundaries as labeled on the right. Lamin A/C, an inner nuclear membrane marker. (B) Fold changes of VE-induced Emerin expression in excitatory and inhibitory neuron proteomics datasets from P56 (adult) and P28 (critical period) mice. Fold changes are calculated as VE versus DK. n=6 biological replicates. One-sample t-test comparing VE/DK fold change to 1. Bars indicate mean and SEM. n.s., not significant. (C) Western blot validation of VE-induced Emerin upregulation in bulk tissue sample from mouse visual cortex. Representative blots are shown at the bottom. n=6, Welch’s t-test. Lines indicate median and 95% CI. (D) Representative immunolabeling of endogenous Emerin protein in cultured mouse neurons. High resolution images of Emerin show that Emerin localizes to both the nuclear envelope and the endoplasmic reticulum at basal level.

**Figure S4.**
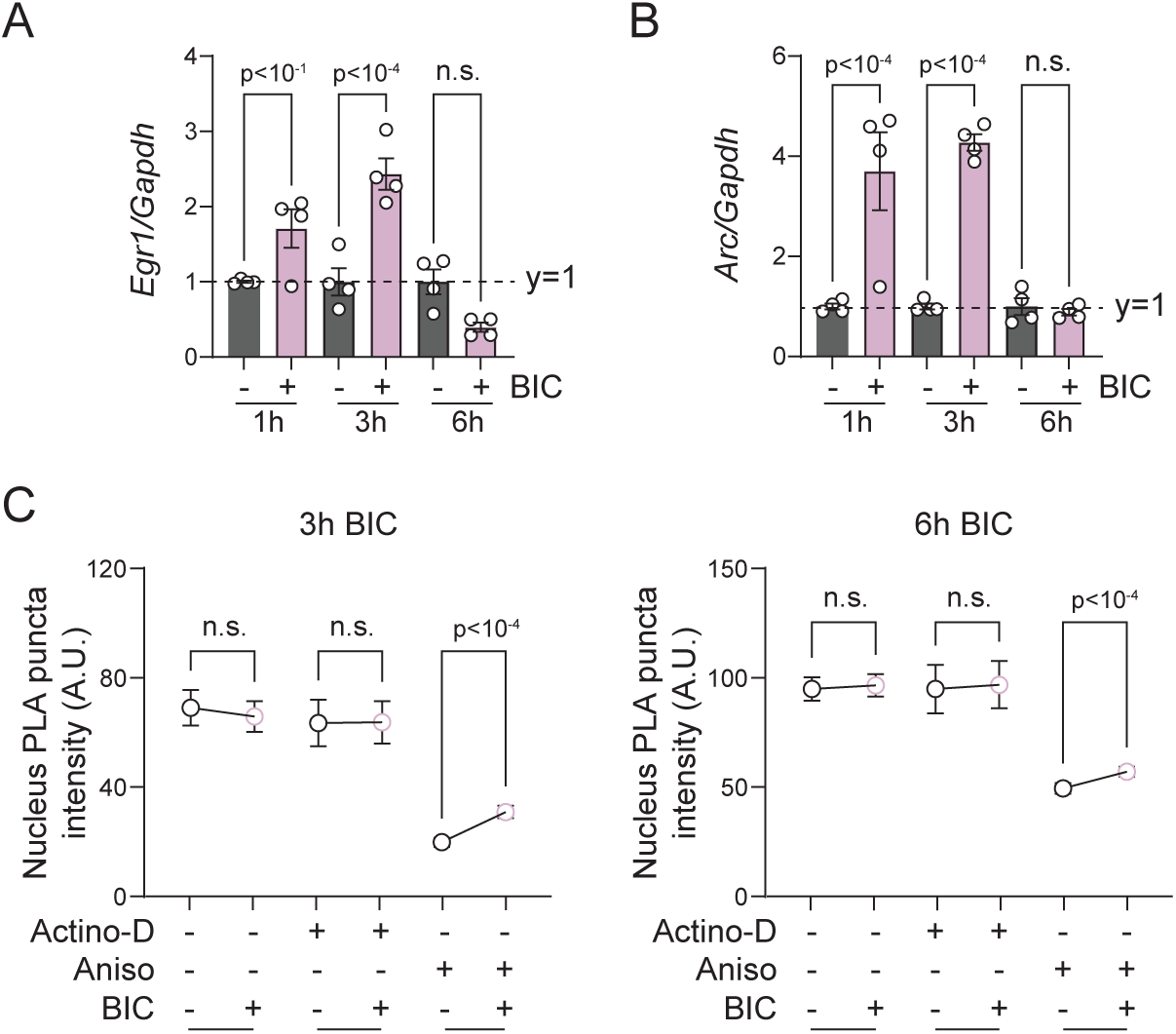
Analysis of bicuculline-induced neuronal activation (related to Figure 3). (A-B) RT-qPCR analysis of immediate early genes *Egr1* (A) and *Arc* (B) to assess neuronal activity-dependent transcriptional induction. All values are normalized to *Gapdh* mRNA level, and then compared to the corresponding controls. n=4 individual wells from 2 independent culture preparations. One-way ANOVA with Sidak multiple comparison (comparing BIC to vehicle at each timepoints). Bars and lines indicate mean and SEM. n.s., not significant. (C) Quantification of nuclear BONCAT-PLA intensity of nascent Emerin at 3h (Left) and 6h (Right) from *in situ* BONCAT-PLA, with pretreatments of transcriptional and translational inhibitors. n=1000-1500 individual neurons from 6 wells of 2 independent culture preparations. Kruskal-Wallis test with Dunn’s multiple comparison (comparing BIC to vehicle for each timepoint and inhibitor treatment). Dots and lines indicate median and 95% CI. n.s., not significant.

**Figure S5.**
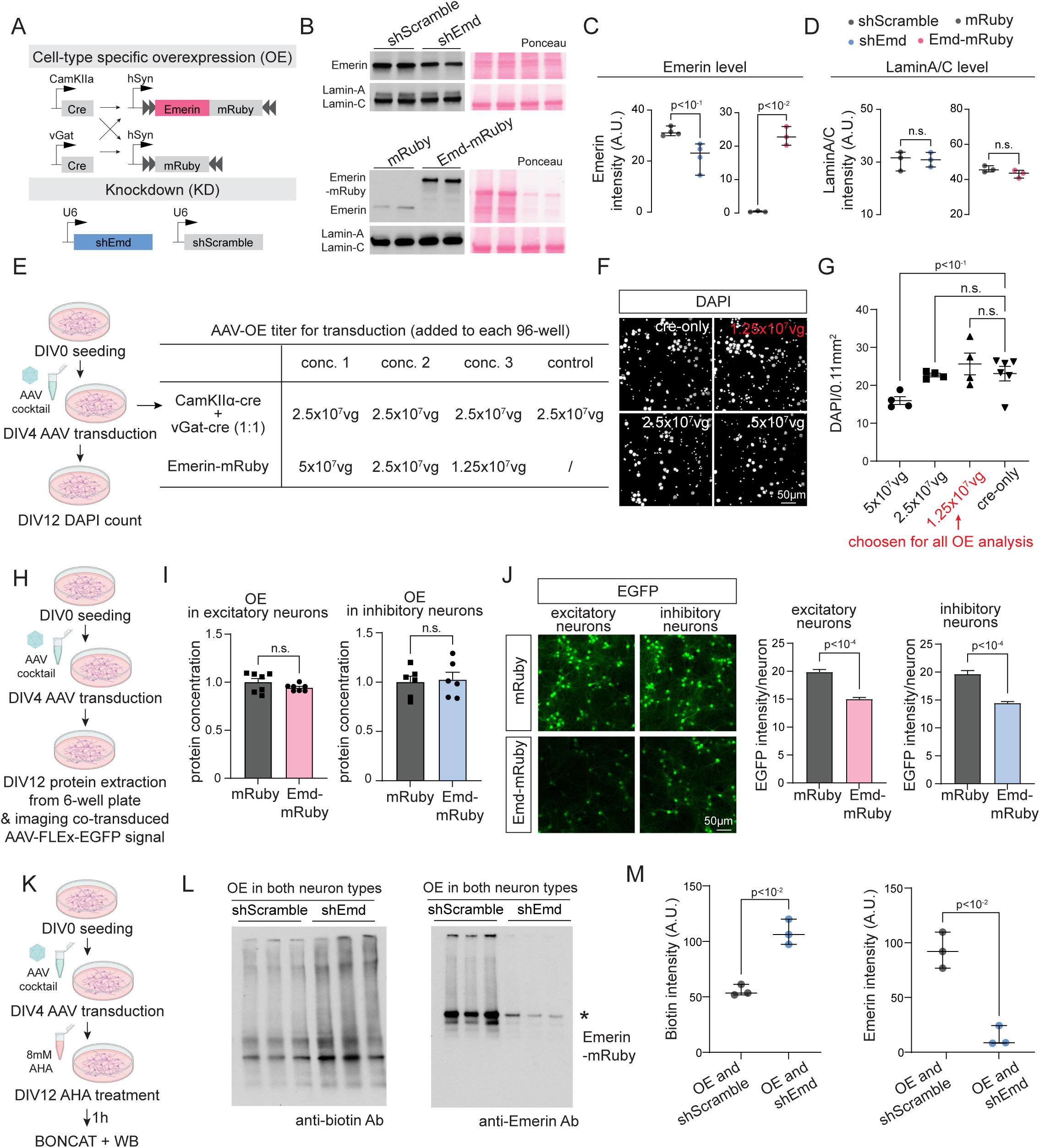
Validation of Emerin KD and OE constructs and the effect of Emerin OE on neuronal density, protein content, and protein synthesis (related to Figure 4). (A-D) Validation of Emerin KD and OE construct in primary neuronal cultures using western blots. (A) Schematics of AAV constructs expressing shEmd, cre-dependent Emd-mRuby, cell type-specific cre recombinase and the corresponding KD and OE controls. (B) Representative WB images probing Emerin levels under Emerin OE and KD conditions. Emerin OE was tested with CamKIIa-cre AAVs. Lamin A/C was also probed as a negative control. When probing Emerin level with Emerin OE, we found that Emerin-mRuby expression level is significantly higher than the endogenous Emerin, making it undetectable without saturating OE signals. To visualize the endogenous Emerin in the control samples and calculate fold change, OE lanes were loaded with 8x less protein. (C) Quantification of Emerin intensity with Emerin KD and OE. (D) Quantification of Lamin-A/C intensity with Emerin KD and OE. Chemiluminescent signals were normalized to ponceau. n=3 or 4 individual wells of 2 independent culture preparations. Welch’s t-test. Lines indicate median and 95% CI. n.s., not significant. (E-G) Analysis of cultured neuron density with various amount of Emerin-OE AAV per well in 96 well plate. (E) Experimental workflow and AAV amount tested. (F) Representative DAPI images of cultures treated with different amounts of AAV. (G) Quantification of DAPI counts in each AAV amount. The second highest dosage that didn’t affect neuronal density at DIV12 was used for subsequent OE experiments (highlighted in red). n=4-6 individual wells of 2 independent culture preparations. Brown-Forsythe and Welch ANOVA test with Dunnett’s T3 multiple comparisons (comparing all dosages to cre-only). Lines indicate mean and SEM. n.s., not significant. (H-I) Analysis of total protein content in cultured neurons with Emerin OE. (H) Experimental workflow. (I) Quantification of total protein extracted per well (data normalized to mRuby controls). n=6-7 individual wells from 3 independent culture preparations. Welch’s t-test. Lines indicate mean and SEM. n.s., not significant. (J) Emerin OE decreased the signal intensity of the co-transfected EGFP in both excitatory and inhibitory neurons. Left, representative images of EGFP signals. Right, quantification of EGFP signal. n=500-1000 individual neurons were quantified for each condition. Kolmogorov-Smirnov test. Bars and lines indicate median and 95% CI. n.s not significant. (K-M) Emerin KD rescues the decreased protein synthesis caused by Emerin OE. (K) Experimental workflow. (L) Left, WB analysis of BONCAT signal with Emerin OE + control KD or Emerin OE + Emerin KD. Right, WB analysis of Emerin expression level with Emerin OE + control KD or Emerin OE + Emerin KD. (M) Left, quantification of biotin intensity. Right, quantification of Emerin intensity. Chemiluminescent signals were normalized to ponceau. n=3 individual wells, Welch’s t-test. Lines indicate median and 95% CI.

**Figure S6.**
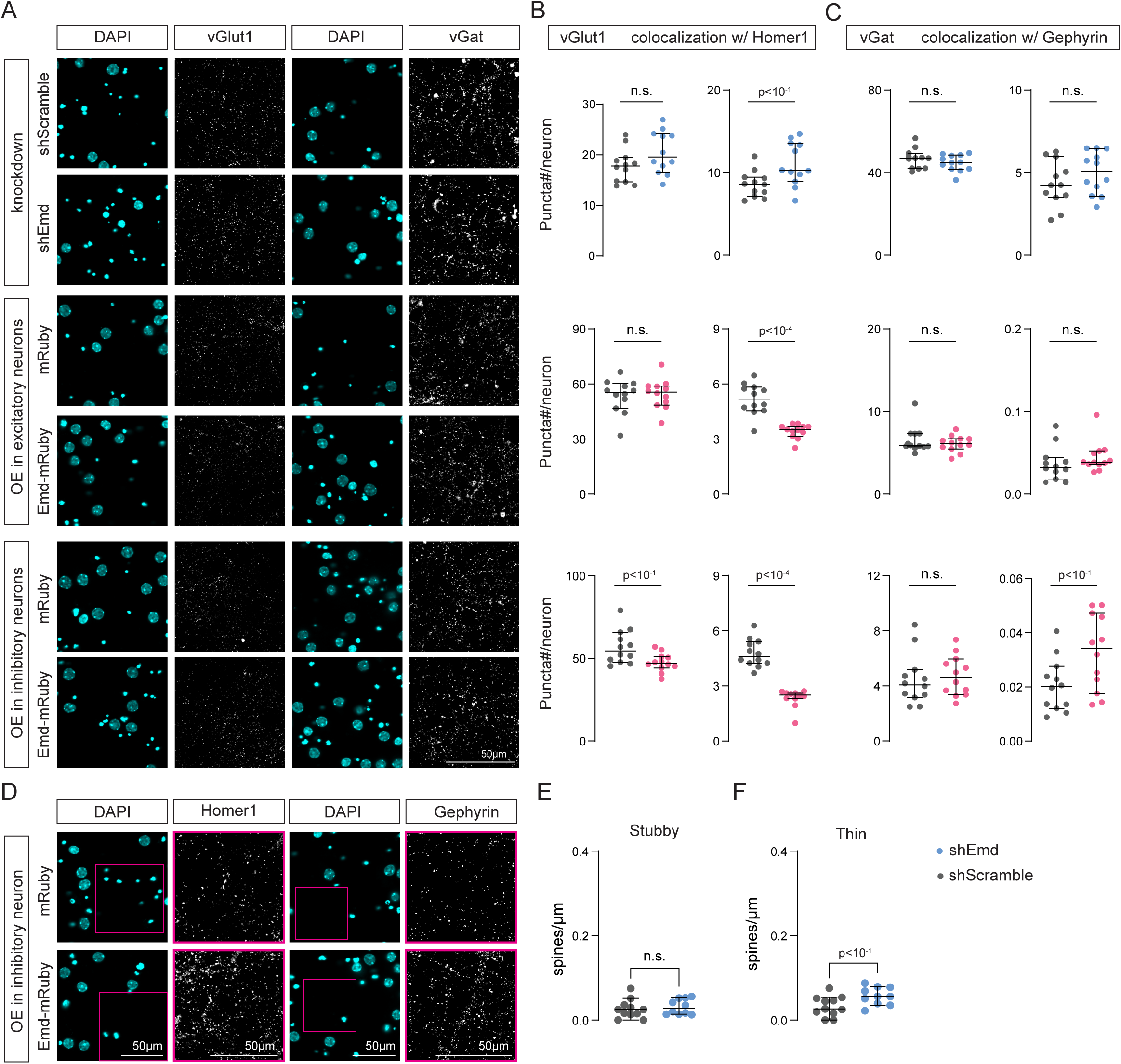
Analysis of synaptic density and spine morphology with Emerin KD and OE (related to Figure 5). (A-D) Analysis of synaptic density with Emerin KD and OE. (A) Representative images of vGlut1 and vGat with Emerin KD (top 2 rows), Emerin OE in excitatory (middle 2 rows) and inhibitory neurons (bottom 2 rows). (B) Quantification of vGlut1 puncta and its colocalization with Homer1 with Emerin KD (top row), Emerin OE in excitatory (middle row) and inhibitory neurons (bottom row). (C) Quantification of vGat puncta and its colocalization with Gephyrin with Emerin KD (top row), Emerin OE in excitatory (middle row) and inhibitory neurons (bottom row). (D) Representative images of Homer1 and Gephyrin with and Emerin OE in inhibitory neurons. Enlarged images of Homer1 and Gephyrin are shown. Puncta counts are normalized to putative neuronal DAPI. n=12 individual wells of 4 independent culture preparations. Mann-Whitney test. Lines indicate median and 95% CI. n.s., not significant. (E-F) Analysis of spine morphology with Emerin KD. Quantifications of stubby spine density (E) and thin spine density (F). n=10-11 dendritic stretches, each from one individual neuron. Data are from 3 independent culture preparations. Mann-Whitney test. Lines indicate median and 95% CI. n.s., not significant.

**Figure S7.**
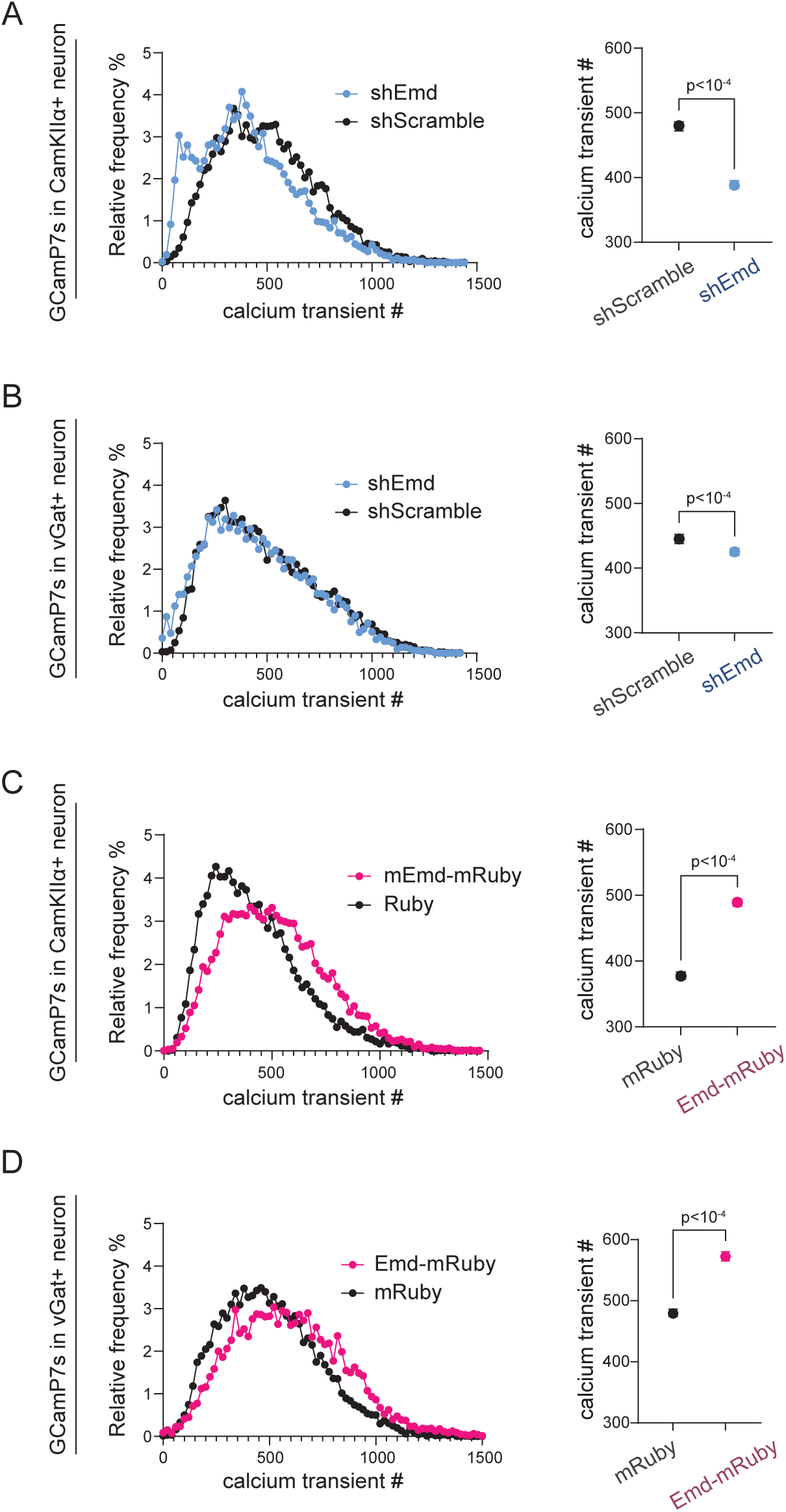
Emerin promotes spontaneous neuronal calcium activity in cultured cortical neurons (related to Figure 6). Spontaneous calcium activity recorded at 20Hz from excitatory neurons with Emerin KD (A), inhibitory neurons with Emerin KD (B), excitatory neurons with Emerin OE (C) and inhibitory neurons with Emerin KD (D). Left, frequency distribution of deconvoluted calcium transient counts over 100s epochs (bin size=20). Right, statistical comparison of calcium transient counts of Emerin KD or OE to their corresponding controls. n=2000-3000 individual neurons from 6 wells of 2 independent culture preparations. Kolmogorov-Smirnov test. Dots and lines indicate median and 95% CI.

**Figure S8.**
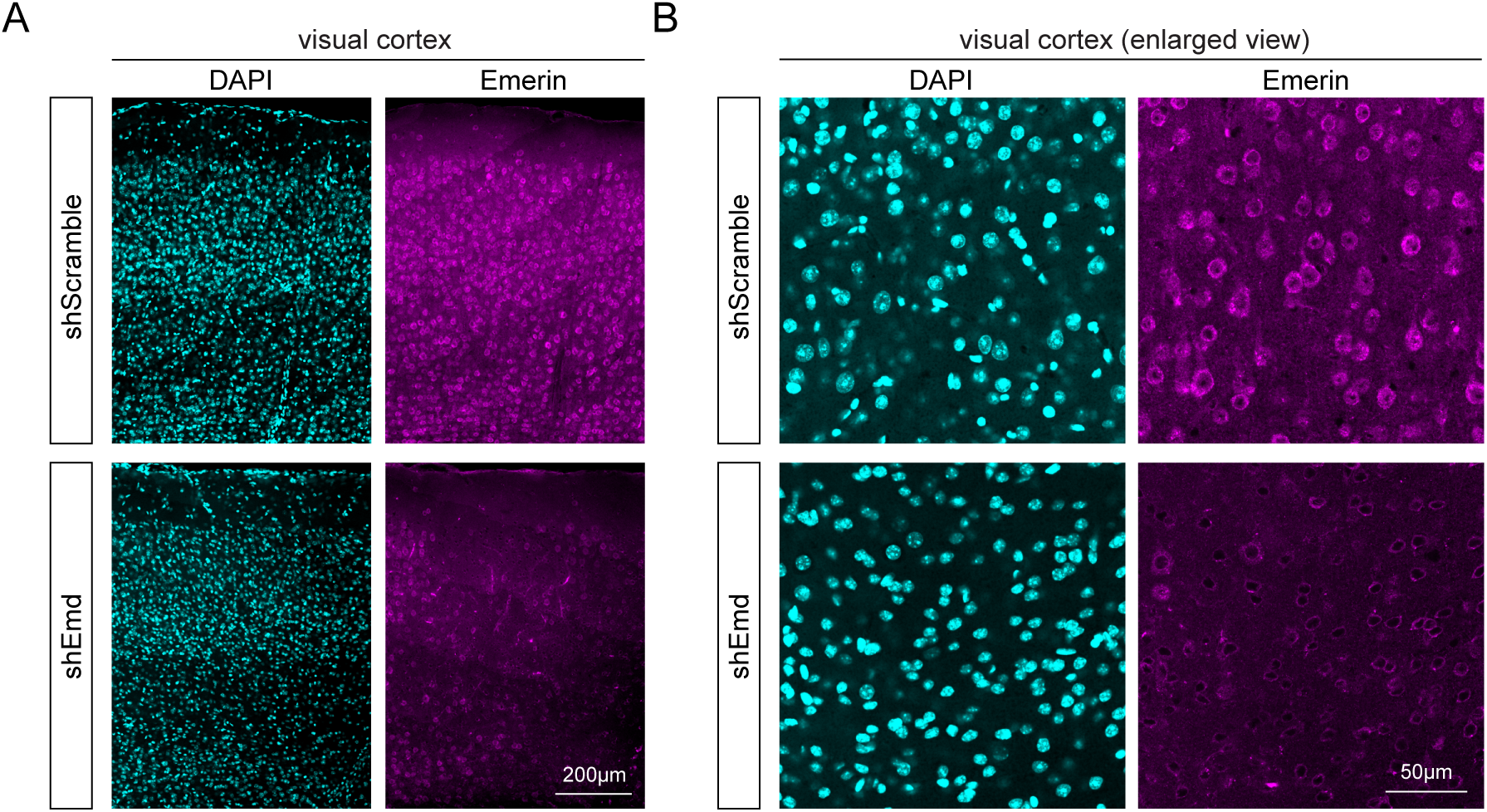
Stereotaxically delivered AAV-shEmd knocks down Emerin expression in mouse visual cortex (related to Figure 6-7) (A) Representative immunolabeling of Emerin protein in the visual cortex of a stereotaxically injected mouse. Left hemisphere received AAV-shEmd injection while right hemisphere received AAV-shScramble injection, which are shown side by side. (B) enlarged view of images in (A)a

## Methods

### Laboratory animals

All animal procedures were conducted with male and female mice according to protocols approved by the Institutional Animal Care and Use Committee (IACUC# 08-0082). Mouse lines EMX1^IRES-Cre^ (JAX 005628), vGat^IRES-Cre^ (JAX 16962) and R26^lsl-GFP-mMetRS^ (JAX 028071) were obtained from Jackson Laboratory. Genotyping was conducted according to protocols published on Jackson Lab websites. To generate homozygous EMX^Cre/WT^-mMetRS^KI/KI^ and vGat^Cre/WT^-mMetRS^KI/KI^ mice, we backcrossed male mMetRS^KI/KI^ to female EMX^Cre/WT^-mMetRS^KI/WT^, and male vGat^Cre/WT^-mMetRS^KI/WT^ to female mMetRS^KI/KI^, respectively, to reduce germline recombination as reported previously [86]. Germline recombination was further tested with PCR for the backcrossed offspring with primers listed below. This primer pair detects the non-recombined R26 insertion allele LSL-GFP-mMetRS in mouse ear tissue, where the forward primer recognizes the LoxP-STOP-LoxP cassette.

Forward: 5’-ctcatgcgttgggtccac-3’

Reverse: 5’-ccggtggtgcagatgaactt-3’

### Primary neuronal cultures from neonatal mice

Primary neurons were cultured with Neurobasal-A media (ThermoFisher, Cat#10888022), supplemented with 2% B27 (ThermoFisher, Cat#17504044), 1% Pen Strep (P/S, ThermoFisher, Cat#15140-122), and 1% GlutaMax (ThermoFisher, Cat#35050061). Complete culture media was pre-warmed to 37°C before use.

P0 mice were euthanized and dissected in 1% P/S-HBSS (HHBS, ThermoFisher, Cat#14175095) to obtain cortical tissue with meninges removed. Cortex tissue was then incubated in digestion buffer (complete culture media with 6% (v/v) papain (Worthington Biochemical, Cat#LS003126) and 750U/µl DNase I (Sigma-Aldrich, Cat# 4716728001)) at 37°C for 20 mins, with periodic agitation. Then, digested tissue was sequentially gravity-precipitated in 20% FBS-HBSS (FBS, Peak Serum, Cat#PS-FB3) and then HBSS. Subsequently, precipitated pellets were gently triturated in dissociation buffer (1x HBSS with 14.44mg/ml MgSO₄ (Sigma-Aldrich, Cat#M2643)). Supernatant was then carefully taken and added to 20% FBS in HBSS and centrifuged 3 mins at 200 rcf. Supernatant was discarded, and cell pellet resuspended in the complete culture media. Neuron density was measured with Countess cell counter (The Countess 3 FL), and neurons were seeded according to the plate format listed below. Multiwell plates were pre-coated with 0.05mg/ml poly-D-lysine (MilliporeSigma, Cat#A-003-E) in Borate buffer (1.25g Boric acid, 1.9g Sodium tetraborate in 400ml ddH_2_O, pH adjusted to 8.5) overnight (O.N.) at 37°C. 1% P/S-HBSS was added to space in between wells to prevent excessive media evaporation. For 96-well plates, wells on the outer rim were not seeded and filled with 1% P/S-HBSS to avoid edge/corner effects.

| Multiwell plate | Experiment | Cat # | Cells seeded/well |
| --- | --- | --- | --- |
| Glass-bottom 96-well culture plate | Imaging | Cellvis LLC, Cat#P96-1.5H-N | 24k |
| Plastic 6-well culture plate | Proteomics/western blot | Greiner, Cat#657160 | 720k |

The cultures were maintained in the complete culture media in a 37°C incubator with 5% CO_2_. At DIV3, FUdR (5-Fluoro-2’-deoxyuridine, 400x stock solution was made with 2mg/ml FUdR (8mM) and 50mg/ml uridine (4mM)) was added to culture media. At DIV4, neurons were transduced with AAVs based on experimental designs. At DIV12, cultured neurons were further treated with labeling, stimulation, or directly harvested for subsequent procedures according to experimental purposes.

### Immunohistochemistry

Animals were anesthetized with isoflurane (Covetrus, Cat#11695067772) and perfused with 1x PBS (ThermoFisher, Cat# 70011044) first, then 4% PFA (Electron Microscopy Sciences, Cat#15710-S) in 1x PBS. Brains were post-fixed in 4% PFA at 4°C O.N., and then switched to 1x PBS. 90µm coronal sections were prepared on a vibratome (Leica VT-1000s). Coronal sections between bregma -2.7mm and -3.64mm were selected for analysis of primary visual cortex. Multiwell dishes containing the sections were wrapped with foil to protect samples from light hereafter, as well as during the whole staining procedure. Sections were stored for up to 4 months at 4°C in 0.5% sodium azide (Sigma-Aldrich, Cat#S2002) in 1x PBS to prevent bacterial growth. All incubation steps hereafter were conducted on a horizontal shaker to provide mild agitation. Sections were incubated in antigen retrieval buffer (100 mM sodium citrate (Sigma-Aldrich, Cat#S-4641), 0.05% Tween-20 (Fisher Bioreagent, Cat#BP337100) in 1x PBS, pH=6.0) for 5 mins at RT, followed by brief boiling, then incubated in antigen retrieval buffer for additional 5 mins at RT. Sections were then washed 3 times in 0.01% PBS-Tx (1xPBS with 0.01% Triton X-100 (Sigma-Aldrich, Cat#X100)) for 10 mins at RT, followed by permeabilization in 0.05% PBS-Tx100 for 1h at RT. Sections were then blocked in IHC blocking buffer (4% bovine serum albumin (BSA, Fisher Bioreagent, Cat#BP1600100), 3% normal donkey serum (Jackson Immuno Research Labs, Cat#017-000-121), 0.1% Triton X-100, and 0.5% sodium azide in 1x PBS) for 1h at RT. Subsequently, sections were incubated with primary antibodies diluted in IHC blocking buffer O.N. at 4°C (see concentrations in Table S6). The next day, sections were washed 3 times with 0.01% PBS-Tx100 for 10 mins at RT, followed by incubation with secondary antibodies diluted in IHC blocking buffer for 2h at RT, then washed 3 times with 0.01% PBS-Tx100 for 10 mins at RT. Finally, sections were mounted on glass slides (Fisher Scientific, Cat#22-034979) with PVA-DABCO (Sigma-Aldrich, catalog #10981) and sealed with a coverslip (VWR, Cat#16004-312). Processed slides were stored at 4°C within a week until imaging.

### Immunocytochemistry

Neuronal cultures in 96-well plates were transferred from the incubator to room temperature (RT). To protect from light, the plates were covered with foil hereafter. Multichannel pipette was used to increase reproducibility of the staining procedure. Pipetting should be as gentle as possible to avoid disturbing the cell layer. Cultures were fixed in 4% PFA in 1x PBS for 20 mins at RT, followed by rinsing in 1x PBS. Cultures were then treated with blocking and permeabilization buffer containing 3% BSA and 0.1% Triton-X 100 in 1x PBS for 30 minutes at RT. After this, cultures were then incubated in primary antibodies diluted to the optimal concentration in ICC blocking buffer (3% bovine serum albumin in 1x PBS) at RT for 2h (see concentrations in Table S6). After a rinse with 1x PBS, cultures were then incubated with the secondary antibodies diluted 1:500 in ICC blocking buffer at RT for 1h. Following a final wash in 1x PBS, the plate was ready for imaging. immunolabeled cultures can be stored at 4°C within a week before imaging.

### Noncanonical amino acid labeling of newly-synthesized proteins

#### in vivo

In mice, 400mM Azidonorleucine (ANL, Chem Impex, Cat#30456) was prepared with UltraPure H_2_O (ThermoFisher, Cat#10977-015) and brought to pH=7 with 10N NaOH (Fisher Scientific, Cat#SS2551). ANL solution was made fresh before treatment and can be kept at 4°C for short-term storage (freeze-thaw sometimes produces hard-to-redissolve precipitation). ANL was administered to mice through i.p. injection at 10 ml/kg using a 31-gauge insulin syringe (Becton Dickinson, Cat#324906) to minimize discomfort. This dosage was tested to be optimal for maximizing proteome labeling without impairing animal health.

#### in vitro

In primary neuronal cultures, 200mM Azidohomoalanine (AHA, Fisher Scientific, Cat#502107772) stock solution was prepared with UltraPure H_2_O and brought to pH=7 with 10N NaOH. Before application, 200mM AHA was diluted in complete culture media pre-warmed to 37°C. The final concentration in each well should be 8mM. For Western blot analysis, cultures were treated with AHA for 1h before harvesting the sample. For proteomic analysis, AHA incubation was extended to 2.5h to maximize signal without affecting neuronal health.

## Dark housing and visual experience

Mice were reared normally under a 12/12 light-dark cycle before being subjected to this protocol. First, mice were dark housed in a light and sound-proof chamber with ventilation and temperature/humidity control for 72h. Mice then received an i.p. injection of ANL, as described above, under dim red light. Then, the DK group was returned to their cage inside the dark housing chamber until tissue collection. The VE group was placed inside a visual experience arena with transparent walls and running wheels, also inside of the dark-housing chamber. 1h after i.p. ANL injection, 6 video tablets were placed to surround the arena, and played a black-white visual noise video to provide visual stimulation including motion and contrast [87]. Freely moving mice were exposed to the visual experience for 4h, and then housed in darkness for another 5h. This 5h period in darkness allows the immediate early wave of visual experience responses to pass in order to capture synthesis and accumulation of newly synthesized effector proteins and maximize signals for proteomic analysis. Tissue collection was then performed under dim red light, where mice were anesthetized with isoflurane and euthanized with cervical dislocation. Brains were then dissected, snap-frozen in isopentane (Millipore Sigma, Cat#270342-2L) incubated on dry ice and stored at -80°C before subsequent processing.

## BONCAT

BONCAT was performed as described previously [88], following the click recipe below and scaled proportionally to protein amount. Proteins were homogenized in DPBS and SDS solution was added to reach 0.05% final concentration to help release lipid-associated proteins. The assembled reaction was then thoroughly vortexed and incubated at RT for 1h. An extra vortex step was performed at 30 min. The click reaction sample was stored at -20°C for up to a week before further analysis.

| Stock solution | per reaction | Cat # | Stock solution recipe |
| --- | --- | --- | --- |
| Protein sample | 500ng | / | / |
| 1.7mM TBTA | 30ul | Sigma-Aldrich,<br>Cat#678937 | 50x stock TBTA: 8.85mg TBTA in 200µl DMSO. 1.7mM TBTA: 20µl 50x TBTA + 180µl DMSO + 800µl tert-butanol. Store at RT |
| 50mM CuSO<sub>4</sub> | 8ul | Sigma-Aldrich,<br>Cat#451657 | 7.98mg/ml in ddH<sub>2</sub>O. Store aliquot in -20°C |
| 5mM Biotin-PEG4-alkyne or heavy/light-labeled-biotin-alkyne | 8ul | Click Chemistry Tools,<br>Cat#TA105-25.<br>Setareh-7889 or -7884 | Dissolve 25mg in 10.927ml DMSO. Store aliquot in -20°C |
| 50mM TCEP | 8ul | Sigma-Aldrich,<br>Cat#C4706 | 14.3mg/ml in ddH<sub>2</sub>O. Make fresh |
| 1x DPBS | top to 400ul | ThermoFisher,<br>Cat#14190-144 | Dilute from 10x DPBS with ddH<sub>2</sub>O |

### Western blot

Cultured primary neurons were extensively washed with 1x DPBS and lysed in the dish with RIPA buffer (G-Biosciences, Cat#786-490) and 1x protease inhibitor (ThermoFisher, Cat#A32963). Dissected brain tissue was lysed and homogenized in 1x DPBS with a hand-held tip sonicator (Fisher Scientific, sonic dismembrator model 100). Protein concentration of the homogenate was determined using either BCA protein assay kit (ThermoFisher, Cat#23225) or DC protein assay kit (Bio-Rad, Cat#5000111). Protein extract with ANL or AHA labeling was clicked with biotin-alkyne described above. 6x Laemmli loading buffer (Boston BioProducts, Cat#BP-111R) was added to protein sample to reach 1x final concentration, vortexed and then boiled for 10 mins at 100°C.

Western blot was conducted using MiniPROTEAN electrophoresis cell and compatible gradient precast gels (Bio-Rad, Cat#456-1096), at 120V constant voltage for 40-50 min. Proteins were transferred to nitrocellulose membranes using Trans-Blot Turbo Transfer System (Bio-Rad, Cat#1704150) and the compatible transfer kit (Bio-Rad, Cat#1704158).

Western blot membranes were subsequently stained with ponceau (Sigma-Aldrich, Cat#09189) for 5 mins at RT. After scanning the ponceau signal on a scanner, the membrane was blocked with 5% blocking reagent (Bio-Rad, Cat#1706404) in 0.05% TBS-Tw20 for 1 h at RT, then incubated with primary antibodies diluted in WB blocking buffer (2.5% bovine serum albumin in 0.1% TBS-Tx100) O.N. at 4°C (see concentrations in Table S6). The next day, the membranes were washed 3 times for 10 mins with 0.05% TBS-Tw20, followed by incubation with secondary antibodies diluted in WB blocking buffer for 2h at RT. Membranes were washed again in 0.05% TBS-Tw20 3 times for 10 mins at RT. Membranes were then incubated with Pierce ECL (ThermoFisher, Cat#32106) for 4 mins and developed onto light-sensitive film (Ece Scientific Co, Cat#E3018) for various amounts of time in a darkroom. Western blot data were analyzed in ImageJ using the lane analysis function.

### Cell type-specific nascent proteomic analysis with DiDBiT in mouse visual cortical tissue

Mouse primary visual cortex was micro-dissected on ice. Specifically, a 3mm-thick section was made from lamda towards anterior using a mouse coronal brain matrices (Ted Pella, Cat#15050). Visual cortex tissue was then homogenized in 200µl 1x DPBS using hand-held tip sonicator. Protein concentration was determined using the BCA protein assay kit.

Visual cortices were dissected and pooled from 5 mice treated under the same condition to accumulate 10mg protein as starting material for subsequent preparation of each MS sample. The following protocol was adapted from DiDBiT and other protocols [38, 88]. Specifically, VE and DK samples were separately clicked with heavy-isotope biotin-alkyne or light-isotope biotin-alkyne, respectively. These two types of biotin-alkyne yield the same click reaction efficiency and do not produce biases, as tested in previous work. Then equal amounts of VE and DK click reaction were mixed, precipitated with methanol-chloroform method, where methanol, chloroform and ddH2O were sequentially added to sample with volume of sample : methanol : chloroform : ddH2O = 1:3:1:3. The mixture was vortexed and centrifuged at 15,000 rcf. Protein precipitates were then resuspended with 8M Urea (MilliporeSigma, Cat#51456) dissolved in 100mM Tris-HCl (pH=8.5). Resuspension was then reduced and alkylated and diluted 4x with 100mM Tris-HCl (pH=8.5). Then TrypZean (MilliporeSigma, Cat#T3568-10mg) was added to protein sample at a 1:20 ratio and incubated on a thermomixer at 37°C. The digested sample was centrifuged at 15,000 rcf for 10min at RT. The supernatant was removed and incubated with 200µl DPBS-washed Neutravidin beads (ThermoFisher, Cat#29200) on a thermomixer for 2h at 22°C. Stringent wash steps were followed as described in the published protocol, with an extra 2M Urea wash right after affinity binding. Bound peptides were eluted with acidic elution buffer (800µl acetonitrile, 197µl UltraPure H_2_O, 2µl formic acid and 1µl trifluoroacetic acid). Eluate was then vacuum-dried with SpeedVac (ThermoFisher, Model#DNA120-115) until all solvent evaporated.

Dried peptides were then resuspended in buffer A (5% acetonitrile, 95% UltraPure H2O and 0.1% formic acid), then analyzed on a Fusion Lumos Orbitrap tribrid mass spectrometer (ThermoFisher). The digest was injected directly onto a 25 cm, 100 µm ID column packed with BEH 1.7 µm C18 resin (Waters). Samples were separated at a flow rate of 400 nl/min on a nLC 1000 (ThermoFisher). Buffer A and B were 0.1% formic acid in 5% and 80% acetonitrile, respectively. A gradient of 1-25% buffer B over 120 min, an increase to 40% buffer B over 40 min, an increase to 90% buffer B over 10 min and held at 90% buffer B for a final 10 min was used for 180 min total run time. The column was re-equilibrated with 20 µl of buffer A prior to the injection of sample. Peptides were eluted directly from the tip of the column and nano-sprayed directly into the mass spectrometer, by application of 2.5 kV voltage at the back of the column. The mass spectrometer was operated in a data-dependent mode. Full MS scans were collected in the Orbitrap at 120K resolution with a mass range of 400 to 1500 m/z and an AGC target of 4e5. The cycle time was set to 3s, and within this 3s the most abundant ions per scan were selected for CID MS/MS in the ion trap with an AGC target of 2e4 and minimum intensity of 5000. Maximum fill times were set to 50 ms and 100 ms for MS and MS/MS scans respectively. Quadrupole isolation at 1.6 m/z was used, monoisotopic precursor selection was enabled, and dynamic exclusion was used with exclusion duration of 5 sec. Proteomics data quantification and statistical analysis were performed on Integrated Proteomics Pipeline (IP2) (Bruker Scientific, http://www.bruker.com).

### Nascent proteomic analysis with DiDBiT on cultured neurons

AHA-labeled cultured neurons were extensively washed with 1x DPBS before lysed in dish with 10% RIPA buffer (diluted in 1x DPBS to reduce the amount of detergent so not to interfere with click chemistry) and 1x protease inhibitor. A cell scraper was used to maximize the sample recovery. Protein concentration was determined with the DC protein assay, and protein samples treated under the same condition were pooled to get 2mg protein starting material for subsequent preparation of one biological replicate. Sample preparation and LC-MS/MS protocol are the same as *in vivo* ANL-labeled samples.

### FUNCAT on cultured neurons

FUNCAT protocols for in vitro and in vivo samples were adopted from previously published guidelines [89] with a few modifications. Specifically, cultured neurons in 96-well plates were transferred from the incubator to RT. From this point on, the plate was wrapped in foil to protect it from light. Cultures were fixed with FUNCAT fix buffer (1mM MgCl2 (Sigma-Aldrich, Cat#M8266), 0.1mM CaCl2 (Sigma-Aldrich, Cat#C3881), 4% PFA, 4% sucrose (Sigma-Aldrich, Cat#G5400) in 1x PBS) for 20 mins at RT, then permeabilized with 0.5% DPBS-Tx100 for 15 mins at RT, followed by blocking with 0.1% Triton X-100 in B-Block (10% normal horse serum (Jackson Immuno Research Labs, Cat#008-000-121), 5% sucrose and 2% BSA in 1x PBS, pH = 7.4) for 1h at RT. Cultures were then washed with 0.1% DPBS-Tx100 for 15 mins at RT. After washing, cultures were incubated in FUNCAT mix O.N. at RT, which was prepared with 2.4µl 85mM TBTA, 50mM TCEP, 2µl 2mM fluor-alkyne (AlexaFluor647-Alkyne (ThermoFisher, Cat#A10278) or AlexaFluor488-Alkyne (ThermoFisher, Cat#A10267)) and 4µl 50mM CuSO_4_ mixed in 1ml fresh 1x DPBS. FUNCAT mix was filtered with a 5ml syringe and 0.22 µm syringe filter (MilliporeSigma, Cat#SLGPR33RS) to remove precipitates, if any. In each 96-well, enough FUNCAT mix was added to fill up the entire well, and the plate was sealed off with adhesive plastic film (Bio-Rad, Cat#MSB1001), and incubated inverted while covered with foil. This prevented adhesion of click reagent aggregates and significantly reduced background aggregates to almost none in our experience. The next day, the film was peeled off and the culture was further washed with 0.1% DPBS-Tx100 for 10 mins at RT and the regular ICC protocol was used to label necessary markers before being imaged.

### Pentylenetetrazol, bicuculline, actinomycin-D and anisomycin treatment

In adult mice, 50mg/kg PTZ (dissolved in 1x PBS) or PBS control was i.p. injected to induce seizure. PTZ solution was made fresh before use. Mice were monitored for 30 minutes after injection to rate seizure level according to the Racine-Pinel-Rovner seizure behavioral scale. Only mice reaching stage 6 or 7 were included in the following analysis. Stage 8 usually quickly transitioned to death, therefore was not included in this study. In total, about 80% of mice injected with PTZ fit the criteria and were included in this study.

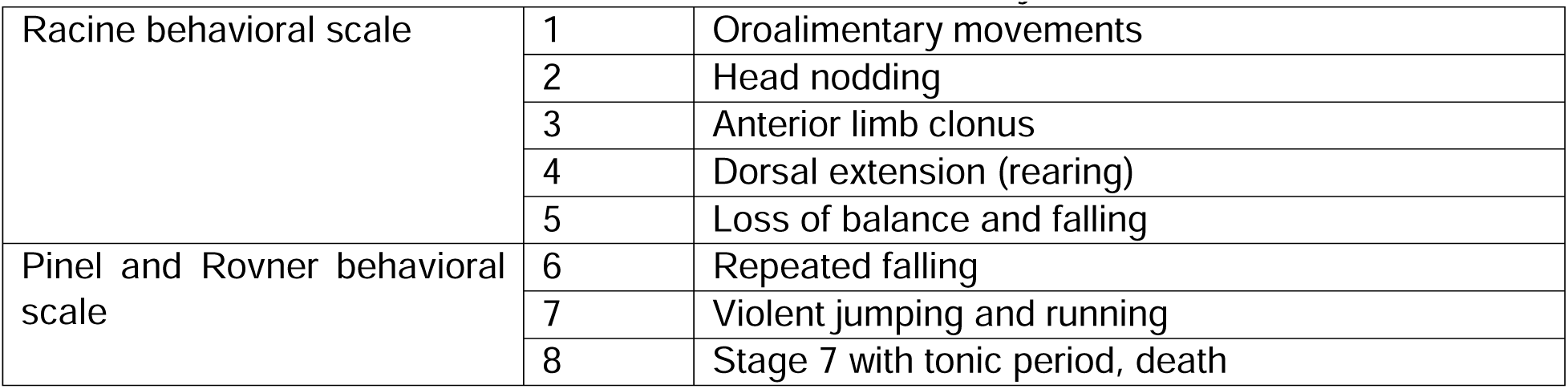

For primary neuronal cultures, 10mM bicuculline (Bicuculline Methiodine, Tocris, Cat# 2503), 200x stock solution was prepared by dissolving powder into ddH_2_O. The stock solution was stored as aliquots in -20°C. When treating neuronal cultures, an appropriate amount of 200x bicuculline stock was mixed with pre-warmed culture media and then added to each well, to reach final concentration of 50 µM in culture. ddH_2_O was used as control for this experiment.

For primary neuronal cultures, 2.5mM actinomycin-D (Sigma Aldrich, Cat#A1410) 100x stock solution and anisomycin (Tocris, Cat#22862-76-6) 1250x stock solution were prepared by dissolving into DMSO (Fisher Scientific, Cat#BP231-100). Stock solutions were stored as aliquots in -20°C. When treating neuronal cultures, appropriate amount of stock solution was mixed with pre-warmed culture media and then added to each well, to reach final concentration of 25µM of actinomycin-D or 40µM of anisomycin in culture. DMSO was used as control for this experiment.

### Reverse transcription qPCR

To perform RT-qPCR on cultured neurons, culture media was removed, and cells were collected with TRIzol reagent (Invitrogen, Cat#15596026), with a cell scraper. RNA microprep kit (Zymo Research, Cat#R1050) was used to extract total RNA from the neuron sample in each well, following the protocol specified by the kit. RNA concentration was measured with nanodrop, and the same amount of RNA was taken from each sample for reverse transcription. cDNA was synthesized using the SuperScript IV reverse transcriptase kit (Invitrogen, Cat#18090010) with random hexamers. qPCR was performed with 2ng/µl cDNA as template, SYBR green master mix (Bio-Rad, Cat#1725121), primers for genes listed below and the Bio-Rad CFX384 qPCR system (Bio-Rad, Cat#1855484). Primers for immediate early genes were obtained from previously published sequences [90].Others were provided by IDT PrimerTime^TM^ qPCR primer assays services.

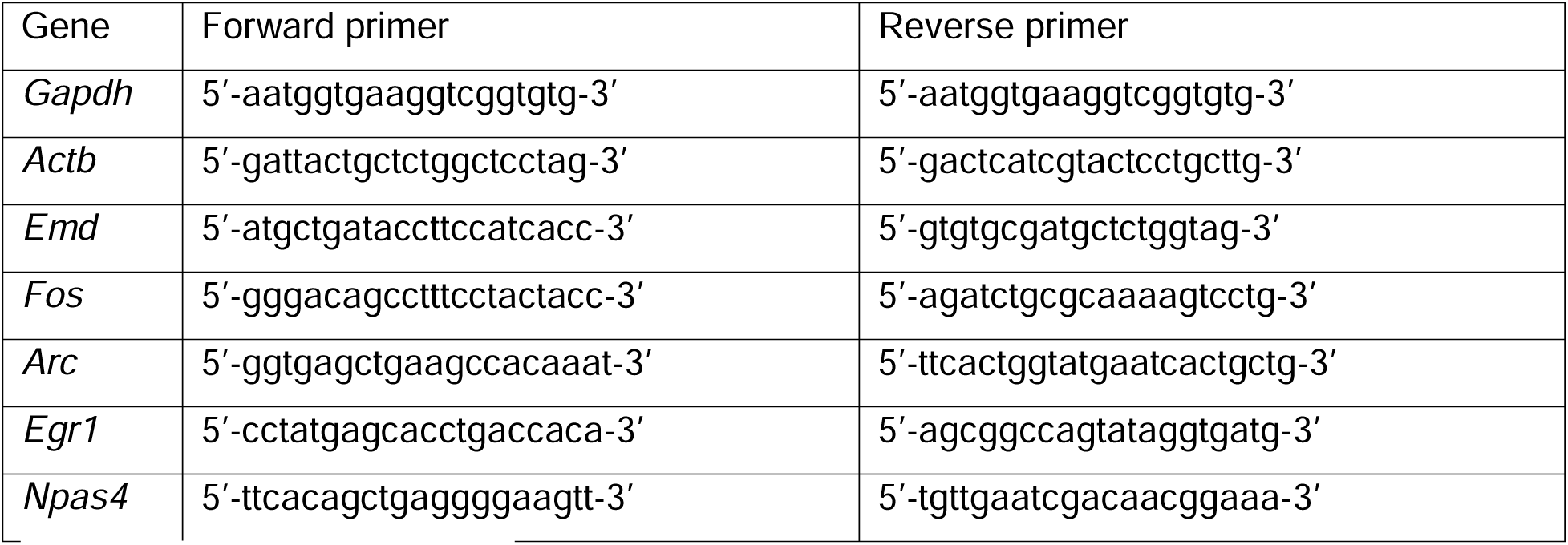

### *In situ* BONCAT-PLA *in vitro*

PLA was performed to detect biotinylated newly-synthesized Emerin. Specifically, *in situ* BONCAT was performed following the *in vitro* FUNCAT protocol, substituting the 2µl 2mM fluor-alkyne with 0.8µl 5mM biotin-alkyne in the FUNCAT mix. The same inverted incubation method was also applied here. After in situ BONCAT, PLA was immediately performed using Duolink® Proximity Ligation Assay kit (Sigma Aldrich) according to the manufacturer’s protocol. Briefly, cultures were blocked with ICC blocking buffer, and then incubated with primary antibodies, goat-anti-biotin and rabbit-anti-Emerin, diluted in ICC blocking buffer, following the ICC protocol of these two steps. Then, the culture was washed with PLA wash buffer A twice for 5 mins at RT, then incubated with PLA probe diluted 1:5 in PLA antibody diluent (Anti-Goat-PLUS (DUO92003) and Anti-Rabbit-MINUS (DUO92005) probes) for 1h in a 37°C humidified oven. Following probe incubation, the culture was rinsed with PLA buffer A twice for 5 mins at RT. Ligation and amplification with far-red detection reagent (DUO92013) were then performed strictly following the manufacturer’s protocol. After the final rinse with 0.01x PLA wash buffer B, the culture was quickly fixed with 4% PFA in PBS for 15 mins at RT to crosslink and secure the PLA complex, and to quickly denature all the reagents used prior to this step. Then, the culture was further subjected to regular ICC protocol to label necessary markers before being imaged.

### Molecular cloning

All plasmids used in this study have an AAV compatible backbone and were subsequently packaged into AAV. A template plasmid AAV-U6-MCS1-hSyn-FLEx-MCS2-mRuby-MCS3 was generated to build shRNA and fusion protein overexpression vectors. Digestion of the multiple cloning site (MCS) was performed using restriction enzyme from NEB Bio-Lab. Three shRNA sequences specifically targeting mouse Emerin mRNA were chosen from The Broad Institute RNAi Consortium shRNA library, with 1 against 3’UTR (TRCN0000240585) and 2 against CDS (TRCN0000216203 and TRCN0000240584). A scramble shRNA sequence was used as control shRNA. DNA oligoes were designed to have sticky ends compatible with MCS1, annealed, phosphorylated and inserted into template vector through T4 ligation (NEB Bio-Lab, Cat#M0202M). For the Emerin-mRuby fusion protein overexpression vector, Emerin coding sequence was PCR amplified from cDNA library reverse-transcribed from mouse primary neuronal culture RNA extract (gift from Maximov lab). PCR primers were designed according to mRNA sequence from NCBI Reference Sequence Database (NM_007927.3). Amplified cDNA was validated through gel electrophoresis and sequencing, then purified and inserted into MCS2 or MCS3 of the template plasmid using the In-Fusion assembly mix (Takara, Cat#638948). The template plasmid was used directly as the overexpression control. Both C-terminus and N-terminus mRuby fusion plasmids were created this way. Ligation/assembly was used to transform NEB Stable competent E. coli (NEB Bio-Lab, Cat#C3040H), spread on carbenicillin (Gibco, Cat#10177012) LB-agar plates. Single colonies were picked, miniprepped and sequenced. Clones with the correct insertion were picked based on the sequencing data, and then subsequently tested with HEK cell transfection. Both C-terminus and N-terminus tagged fusion protein showed ER and nucleus membrane localization, same as the endogenous Emerin localization pattern stained with antibody in mRuby-only control transfected HEK cell. The C-terminus fusion protein is expressed at a higher level than the N-terminus fusion protein. Therefore, the C-terminus fusion plasmid was used for AAV packaging and future experiments. shEmd triple-transfection in HEK cells overexpressing mouse Emerin showed efficient mouse Emerin knockdown, compared to sham transfected as well as shScramble transfected. AAV-CaMKIIα-Cre was modified from AAV-CaMKIIα-EGFP-Cre from Addgene (Addgene #105551) by removing the EGFP sequence. AAV-vGat-cre was made by replacing the promotor in AAV-CaMKIIα-Cre with the vGat promotor from hU6-gRNA-vGat-dCas9-EGFP described before [91]. AAV-hSyn-FLEx-jGCaMP7s was obtained from Addgene (Addgene #104491).

### AAV production

AAV packaging was performed as previously described [92]. First, plasmids to be packaged were maxi-prepped (Zymo Research, Cat#D4203) according to the protocol described by the kit. AAV-293 HEK cells (Agilent, Cat#240073) were used and cultured in 15cm culture dishes with 5% FBS-DMEM (ThermoFisher, Cat#11995065) supplemented with 1% NEAA (ThermoFisher, Cat#11140-050), 1% Sodium pyruvate (ThermoFisher, Cat#11360-070), 1% GlutaMax and 1% P/S. Briefly, HEK cells were triple-transfected with pHelper, pAAV.DJ and the expression vector using PEI (Polysciences, Cat#23966-1). Culture media was collected 72h and 120h after transfection. Cells are scrapped and collected at the 120h time point. Viral particles are precipitated from media with PEG. The media pellet and the cell pellet were digested separately using salt activated enzyme and then combined after completion. Ultrahigh-speed centrifugation was used to purify viral particles in a density gradient using Optiprep. The purified virus was then subjected to 4 rounds of DPBS buffer exchange in Amicon filter tubes and concentrated to the desired volume during the final buffer exchange. AAV were titered using the qPCR protocol described in the paper cited above. From 5 15cm culture dishes, we usually recovered virus with a titer around 10^13^ vg/ml in 400-600µl final volume. The virus was then aliquoted and stored in -80C, or 4C after thawing. For primary culture experiments, AAV-GCaMP and AAV-shRNA were added to primary culture at a dosage of 5x10^7^vg per 96 well, and cre-delivering AAVs

2.5x10^7^vg per 96 well. Emerin OE AAVs and the corresponding control were added at 1.25x10^7^vg per 96 well, with 1.25x10^7^vg cre-delivering AAVs. In other well plate formats, virus dosage was adjusted proportionally to the surface area. This dosage allowed sufficient fluorescent signal for imaging and good OE and KD level for Emerin in our experiments, while maintaining good culture health. For stereotaxic injections, viruses were used at a concentration of 1x10^12^vg/ml.

### Calcium imaging *in vitro*

For calcium imaging in vitro, DIV4 primary mouse neurons were transduced with AAV-hSyn-FLEx-GCaMP7s and either AAV-CaMKIIα-cre or AAV-vGat-Cre. AAVs reach steady expression 7 days after transduction. Neuronal cultures were imaged on DIV12. Calcium transients were imaged using the ImageExpress HT.ai confocal high-content imaging system (IXM, Molecular Devices). Glass bottom 96 well plates were placed in the imaging system with environmental control (37C, humidified gas constituted by 5% CO2, 21% O2 and rest with N2) throughout the entire imaging period. A 10x Plan Apo 0.45NA objective was used with a confocal 50µm slit setting. Laser power is set at 2% and each frame is exposed for 50ms, equal to 20Hz sampling rate. Each well was imaged sequentially in 4 non-overlapping FOV, with each FOV was imaged for 2min. The area imaged in total accounts for 25.45% of the well area. Single z-plane single-channel t-series .TIF files were analyzed using Suite2P [93]. Default settings were used for ROI registration and spike detection. ROI detection was manually inspected for each movie file as a quality control. For each movie file, the total deconvoluted spike counts and averaged spike amplitude were calculated for each ROI using Python and Numpy package. Data visualization and statistics were performed using Prism GraphPad.

### Stereotaxic injection in visual cortex

To deliver AAV stereotaxically into the binocular region of adult mouse visual cortex, mice were anesthetized with 5% isoflurane inside an anesthesia induction chamber, and then head-fixed on stereotaxic frame (Kopf instruments, model-942 with digital display) with 1.5-2% isoflurane constantly delivered through a nose cone throughout the procedure. 0.25mg/ml Flunixin (supplied by TSRI department of animal resources) was i.p. injected at 10ml/kg and 0.3mg/ml buprenorphine (Covetrus, Cat#055175) was i.m. injected at 2ml/kg prior to surgery procedures for analgesic purposes. The hair was removed, and the exposed skin was sterilized with Povidone-Iodine as well as 75% ethanol. A midline incision was made to expose the skull above the visual cortex. A NanoFil 10µl syringe (WPI, Cat#NANOFIL) with 33-gauge beveled NanoFil needle (WPI, Cat#NF33BV-2) was loaded with AAV, motorized with an electrical micropump (WPI, Cat#UMP3T-2) to control the amount of liquid dispensed and navigated to visual cortex through the digital coordinates display. Bregma was used as set to (0, 0, 0) reference. Coordinates for the binocular region of primary visual cortex are AP = -3.3, ML = +/- 2.8, and the needle was advanced along the dorsal-ventral axis with 2 stops at DV = -1.1 and -1.5 for virus injection. A small opening was made in the skull with microdrill (RWD, Cat#78001) above the site selected for injection. The syringe was advanced slowly to the first injection site. At each DV stop, 600nl of AAV was dispensed at 80nl/min. The syringe remained at each injection site for another 10min to increase local AAV diffusion. After injection and retraction of the needle, the skin was sutured (Healthcare Products, Cat#ETHJ392H), and the closure was further reinforced with Vetbond. Mice continued to receive the same analgesics regime for 3 days and were monitored for health daily. Animals were used for experiments after 4 weeks to allow full expression of viral proteins.

### Visually evoked potential surgery and recording

For VEP recording experiments, we used the Data Acquisition and Analysis MP150 System (BIOPAC Systems, Inc.) to aid in implanting the electrode and for VEP recording. AAV was delivered through the stereotaxic injection protocol as described above. After AAV was injected, 2 slightly larger openings were drilled at coordinates AP = -3.5, ML = +/- 3.2 to implant the recording electrodes bilaterally into the binocular regions of visual cortex, as described previously [94]. 2 additional openings were drilled at coordinates AP = +1, ML = +/-1 for bilaterally implanting reference electrodes in the prefrontal cortex. The electrodes consist of an insulated stainless-steel wire (0.005” diameter, 3 mm length, 005SW-30S, PlasticsOne Inc.). The insulation was burned to remove 2 mm of coating from one end of the wire and then it was inserted into a male pin (A-M systems, Protech International Inc) and crimped (AFM8 crimping tool, Bloomfield Hills, Mich). The insulation at the other end of the wire was burned to remove 0.5 mm of coating with the help of a magnifying glass and a ruler. The male pin was connected to a female miniature pin (0.032”x0.4”0.005” diameter, 3 mm length, 005SW-30S, PlasticsOne Inc.), crimped to a standard touchproof connector to interfacing to 100C-series Biopotential amplifiers (Biopac Inc). The average active electrode impedance was 5.2 ± 2.42 kΩ (mean ± SD, Grass Electrode Impedance Meter, Model EZM3A, Quincy, MA).

Two reference electrodes were implanted first, slowly inserted 0.25mm under the dura, and secured in place with dental cement (Stoeling, Cat#100229-622). Two recording electrodes were implanted next, one by one. During the implantation, the recording electrode was held on a stereotaxic arm and connected to the MP150 amplifier system with real-time recording with AcqKnowledge data acquisition software. The corresponding reference electrode was connected during this process to test the circuit. As the recording electrode was lowered into the brain, the noise level significantly dropped when it touched the dura, and from this point, the recording electrode was further advanced for 250µm to reach layer 2/3 of visual cortex. The recording electrodes were then cemented to the skull with dental cement. Then, a 3-pronged head bar (design kindly shared by the Komiyama lab, UCSD) was cemented to the skull, providing head restraint during the VEP recording with a mini-clamp designed to fit the 3-pronged head bar. Mice were post-operatively cared for in the same way as the stereotaxic injection protocol.

4 weeks after implanting the electrodes, VEP recording was performed under dim red light at 4-6pm period with the mice dark-adapted for 1h prior to each session. Mice were anesthetized with 10mg/ml xylazine (AnaSed, Cat#59399-110-20) i.p. injected at 3ml/kg and 10mg/ml pentobarbital (Pisa agropecuaria, PISAPENTAL) i.m. injected at 3ml/kg. Mice were placed in the head-restraint mini-clamp and positioned with their eyes 20cm in front of a LED stroboscope (Monarch instruments, Nova-pro 300) controlled by the MP150 system to flash at 1Hz of 0.05 cd s/m^2^ white light. 250 flashes were delivered binocularly while simultaneously recording visually evoked response from both hemispheres. The signal was amplified 10,000 (PM150 system, Biopac Inc), band pass filtered (1–35 Hz) and digitized at 20000 Hz with 12 bits resolution. Responses were averaged across 250 iterations within 1 session. 2 sets of 2 amplifiers were used to measure each hemisphere, and then switched between the 2 hemispheres and recorded again to exclude potential instrument bias. 4 traces were obtained for each visual cortex response and averaged to quantify response amplitude and latency.

### Visual cliff test and video analysis

The visual cliff apparatus is a home-made acrylic plexiglass box as depicted in Figure 7, placed inside a dark room with dim red light. The top part is a 60cm x 60cm x 60cm plexiglass box enclosing a flat transparent floor with black and white checkerboard pattern at 100% contrast attached to the underside of one quadrant of the floor (the safe zone). White paper was taped around the outside to block irrelevant visual cues. The bottom part of the apparatus is 60cm x 60cm x 50cm in size, with the same black-white checkerboard pattern covering the 4 walls and the floor. A Logitech Brio 4K webcam was held right over the top of the arena walls recording at 30 frames per second. A light source was placed next to the webcam to provide lighting for the test area.

With this setup, the texture of the safe zone and the remainder of the transparent floor on the top part is homogenous to the mice, and the proximal visual cues inside the safe zone can be distinguished from the distant visual cues on the lower floor of the unsafe zones. Our visual cliff test, slightly different from the conventional visual cliff setup, is constituted by 25% of safe zone and 75% unsafe zone, as opposed to the traditional 50-50 division. This setup doubles the chances of crossing the edge from safe to unsafe zones (2 edges as opposed to 1 edge in 50-50 division), which seemed to help animals recognize the differences between the safe and unsafe zones sooner and decide to stay in the safe zone faster if their depth perception was intact. All visual cliff tests were performed at 1-3pm. Prior to tests, mice were acclimated in the testing room for 1h. During the test, a mouse was placed in the center of the safe zone, and then allowed to explore freely for a 2min period. Apparent safe zone preference was detected within this 2min period, and this preference didn’t change when tested a second time 4 weeks later. The plexiglass surface of the top part (the walls and the floor) was thoroughly cleaned with disinfecting and deodorizing spray (PeroxiGard, AHP, Cat#74559-4) in between trials. Videos were post-processed with Capcut movie processing software to remove the frames involving the experimenters. Then, mice movements and arena landmarks were labeled and traced using DeepLabCut Resnet50 machine learning network following previously published guidance [95]. Python-based DLCAnalyzer was modified and applied to define safe/unsafe zones, calculate speed/time/distance and generate zone visit reports [96].

### IXM High-content imaging, image processing and analysis

Neuronal cultures in glass-bottom 96 well plates (processed according to the ICC, FUNCAT or *in situ* BONCAT-PLA protocols) were imaged using the HT.ai confocal high-content imaging system (Molecular Devices). The imaging was set up using either the 20x Apo LWD (NA 0.95) or the 40x Apo LWD (NA 1.15) water immersion objectives, with 4 available channels including DAPI, FITC, TRITC and Cy5. 16-36 non-overlapping FOV were imaged per well. All datasets were manually inspected for out-of-focus images or images including well edges as quality control. Images were pre-processed using ImageJ in a batch mode before Cell Profiler analysis [97]. Then, customized Cell Profiler pipelines were written and applied to each dataset based on the experimental designs. In general, a pipeline includes illumination correction, primary ROI identification with threshold and watershed, relating objects, image/object-based intensity measurements, ROI overlay output, and data exportation in database format. Databased files were further read by DB browser software and per-image and/or per-object information were exported to .csv format for statistics.

### Nikon confocal imaging and analysis

Mouse brain sections were imaged on a Nikon A1 confocal microscope with either the 10x Plan Apo objective (NA 0.45), the 20x Plan Apo objective (NA 0.7) or the 40x Plan Fluor oil immersion objective (NA 1.3). cFos expression was analyzed through manual cell counting assisted with ImageJ cell counter plugin.

### Bioinformatics

Bioinformatic analyses were performed using the QIAGEN IPA (QIAGEN Inc.) [98], Metascape database [99] and STRING PPI network analysis [100]. For IPA, core analysis was performed using p-value (0.05 cutoff) and log_2_ expression ratio as data input. The same datasets were submitted to Metascape for parallel analysis. Pathway components identified by Metascape were then input to the STRING database to generate PPI networks with orphan nodes removed.

### Statistics

All statistics were performed using Prism GraphPad. The statistical tests and plot types were specified in the corresponding figure legends. In some plots, confidence intervals or error bars are obscured by the median or the mean. Normalization was noted in the y-axis label or figure legends.

## Notes

### Competing Interest Statement

The authors have declared no competing interest.

